# Cortical resonance selects coherent input

**DOI:** 10.1101/2020.12.09.417782

**Authors:** Christopher Murphy Lewis, Jianguang Ni, Thomas Wunderle, Patrick Jendritza, Andreea Lazar, Ilka Diester, Pascal Fries

## Abstract

Synchronization has been implicated in neuronal communication, but causal evidence remains indirect. We used optogenetics to generate depolarizing currents in pyramidal neurons of cat visual cortex, emulating excitatory synaptic inputs under precise temporal control, while measuring spike output. Cortex transformed constant excitation into strong gamma-band synchronization, revealing the well-known cortical resonance. Increasing excitation with ramps increased the strength and frequency of synchronization. Slow, symmetric excitation profiles revealed hysteresis of power and frequency. Crucially, white-noise input sequences enabled causal analysis of network transmission, establishing that cortical resonance selectively transmits coherent input components. Models composed of recurrently coupled excitatory and inhibitory units uncovered a crucial role of feedback inhibition and suggest that hysteresis can arise through spike-frequency adaptation. The presented approach provides a powerful means to investigate the resonance properties of local circuits and probe how these properties transform input and shape transmission.

## Introduction

The brain’s computational abilities arise from communication within and between neuronal groups, and the dynamic modulation of neuronal communication is believed to enable flexible behavior (Engel et al., 2001; Fries, 2015; Varela et al., 2001). A compelling means to modulate neuronal communication is synchronization (Akam and Kullmann, 2010; Azouz and Gray, 2003; Börgers and Kopell, 2008; Hahn et al., 2014; Palmigiano et al., 2017; Salinas and Sejnowski, 2001; Wang, 2010). Neuronal synchronization is determined by cellular and network properties that define intrinsic timescales for activity. The intrinsic timescale of cells and circuits can be characterized by resonance, i.e. how inputs are amplified, or preferentially transmitted. In single neurons, specific combinations of diverse conductances can establish membrane and firing-rate resonances (Fellous et al., 2001; Hutcheon and Yarom, 2000; Lampl and Yarom, 1997; Schreiber et al., 2004). In networks, interactions between recurrently coupled excitatory and inhibitory neurons generate resonances based on connectivity (Börgers and Kopell, 2003; Buzsáki and Wang, 2012; Tiesinga and Sejnowski, 2009; Whittington and Traub, 2003).

A prominent cortical resonance occurs in the gamma-band (30-90 Hz) (Adesnik and Scanziani, 2010; Cardin et al., 2009; Etter et al., 2019; Iaccarino et al., 2016; Lu et al., 2015; Ni et al., 2016; Sohal et al., 2009). The Communication-through-Coherence (CTC) hypothesis (Fries, 2005, 2015) proposes that gamma-band synchronization between neuronal groups can flexibly determine their communication. Computational models have demonstrated that gamma-rhythmic inputs can entrain a postsynaptic population of recurrently coupled excitatory and inhibitory (E-I) units, thereby enhancing the impact of the entraining input, and reducing the impact of competing inputs (Börgers and Kopell, 2008; Hahn et al., 2014; Palmigiano et al., 2017). This proposal has accrued considerable correlative evidence, for example, gamma-rhythmic gain-modulation of neuronal and behavioral responses (Ni et al., 2016); phase-dependent power-covariation and transfer entropy between neuronal groups (Besserve et al., 2015; Womelsdorf et al., 2007); and selective enhancement of interareal gamma-band synchronization by attention (Bosman et al., 2012; Grothe et al., 2012), which improves behavioral performance (Rohenkohl et al., 2018). However, it has remained difficult to provide direct causal evidence for selective transmission of coherent inputs via network resonance.

Direct evidence for a causal role of synchronization in neuronal communication can be obtained through experimental control of network input and simultaneous measurement of spike output (Akam et al., 2012). We emulated excitatory synaptic input to a local population with millisecond temporal precision using Channelrhodopsin-2 (ChR2), a light-activated cation channel (Boyden et al., 2005). We transfected pyramidal cells in cat visual cortex, a classical model for investigating cortical information processing (Douglas and Martin, 2004). Illumination of ChR2-expressing neurons enabled control of synchronous excitation in vivo.

Stimulation with constant light confirmed the previous finding that cortical networks can transform temporally flat excitatory input into gamma-rhythmic spike output (Adesnik and Scanziani, 2010; Lu et al., 2015; Ni et al., 2016) with features similar to that generated by visual stimulation (Fries et al., 1997; Fries et al., 2002; Gray et al., 1989; Gray and Viana Di Prisco, 1997). Slowly varying the excitation to the network with ramps and symmetric stimulation profiles revealed that the peak frequency of the gamma resonance could vary between 30 and 70 Hz, and that there was pronounced hysteresis for both the power and the frequency. Sinusoidal stimulation demonstrated that network spike output was entrained by rhythmic input with a fidelity that increased up to 40 Hz and decreased slightly for 80 Hz.

Finally, we sought to determine if the intrinsic resonance of visual cortical populations can act as a filter to select coherent components of external excitatory drive. Direct stimulation of excitatory cells with temporal white noise dramatically illustrated that the resonant properties of the local circuit established an endogenous temporal receptive field, or window of opportunity, for external excitatory drive. In contrast to periodic signals (like sinusoids or rhythmic pulse-trains), white noise is not auto-correlated, and therefore enables a causal analysis of network transmission, i.e. from excitatory input to spike output (Bryant and Segundo, 1976; Mainen and Sejnowski, 1995; Marmarelis and Naka, 1972). Spike-triggered averaging of the white-noise light sequence revealed that spikes were preceded by episodes of gamma-rhythmic input. Correspondingly, an analysis of Granger causality between the white noise input and neuronal spike output revealed a pronounced gamma-band peak. Simulations with a well-established recurrent network composed of conductance-based model neurons (Börgers, 2017) reproduced our core results. Modeling confirmed the central role of strong, fast feedback inhibition in gamma-band resonance (Börgers and Kopell, 2003; Sohal et al., 2009; Stark et al., 2014). The essential resonance phenomena were also evident in a greatly simplified network of leak-integrate and fire units. Modeling of the power and frequency hysteresis effects required the addition of a non-inactivating potassium current, the M-current, to the excitatory units. Overall our results suggest that recurrent excitatory-inhibitory coupling establishes intrinsic temporal scales for neuronal activity in local circuits. These intrinsic scales are apparent in the resonant properties of the population, which temporally transform excitatory input, selecting components of time-varying input coherent with the resonant oscillation and attenuating non-coherent components.

## Results

### AAV1 and AAV9 transfect excitatory neurons in cat visual cortex, and constant optogenetic stimulation reveals gamma-band resonance

Recombinant adeno-associated virus (AAV) vectors are widely used as gene-delivery tools (Vasileva and Jessberger, 2005). AAV-mediated expression of Channelrhodopsin-2 (ChR2) has been used in several mammalian species, including mice, rats and non-human primates (Diester et al., 2011; Gerits et al., 2015; Scheyltjens et al., 2015). In this study, three pseudo-typed AAVs, AAV1, AAV5 and AAV9, were applied in visual cortex of the domestic cat (*felis catus*). We injected AAVs carrying the gene for hChR2(H134R)-eYFP under the control of the Ca2+//calmodulin-dependent protein kinase type II alpha (CamKIIα) promoter. Injections targeted either area 17, the cat homologue of primate area V1, or area 21a, the cat homologue of primate area V4 (Payne, 1993). All AAV1 and AAV9 injections resulted in robust transfection (which was not the case for AAV5, see Methods). Transfection was evident in confocal fluorescence microscopy (and often in epifluorescence) and in the neuronal responses evoked by light. In total, we transfected neurons in area 17 in four hemispheres of three cats, and in area 21a in four hemispheres of four cats.

In two cats, after electrophysiological recordings were completed, brains were histologically processed, and slices were stained for parvalbumin (PV) and/or gamma-Aminobutyric acid (GABA) (Fig. 1 and S1). One cat had been injected with AAV1-CamKIIα-hChR2(H134R)-eYFP into area 17. Across several slices and imaging windows of area 17, we identified 264 unequivocally labeled neurons, which showed ChR2-eYFP expression or GABA-anti-body staining; of those, 146 were positive for GABA, and 118 expressed ChR2-eYFP, and there was zero overlap between these groups (Fig. 1A-D). In the same cat, across several additional slices and imaging windows of area 17, we identified 284 unequivocally labeled neurons, which showed ChR2-eYFP expression or PV-anti-body staining; of those, 145 were positive for PV, and 139 expressed ChR2-eYFP, with four neurons showing clear ChR2-eYFP fluorescence and partial (patchy) PV staining (Fig. S1A-D). The other cat had been injected with AAV9-CamKIIα-ChR2-eYFP into area 21a. Across several slices and imaging windows of area 21a, we identified 182 unequivocally labeled neurons, which showed ChR2-eYFP expression or PV-anti-body staining; of those, 73 were positive for PV, and 109 expressed ChR2-eYFP, and there was zero overlap between these groups (Fig. S1E-H). Thus, ChR2 expression occurred almost exclusively in excitatory neurons.

**Figure 1.**
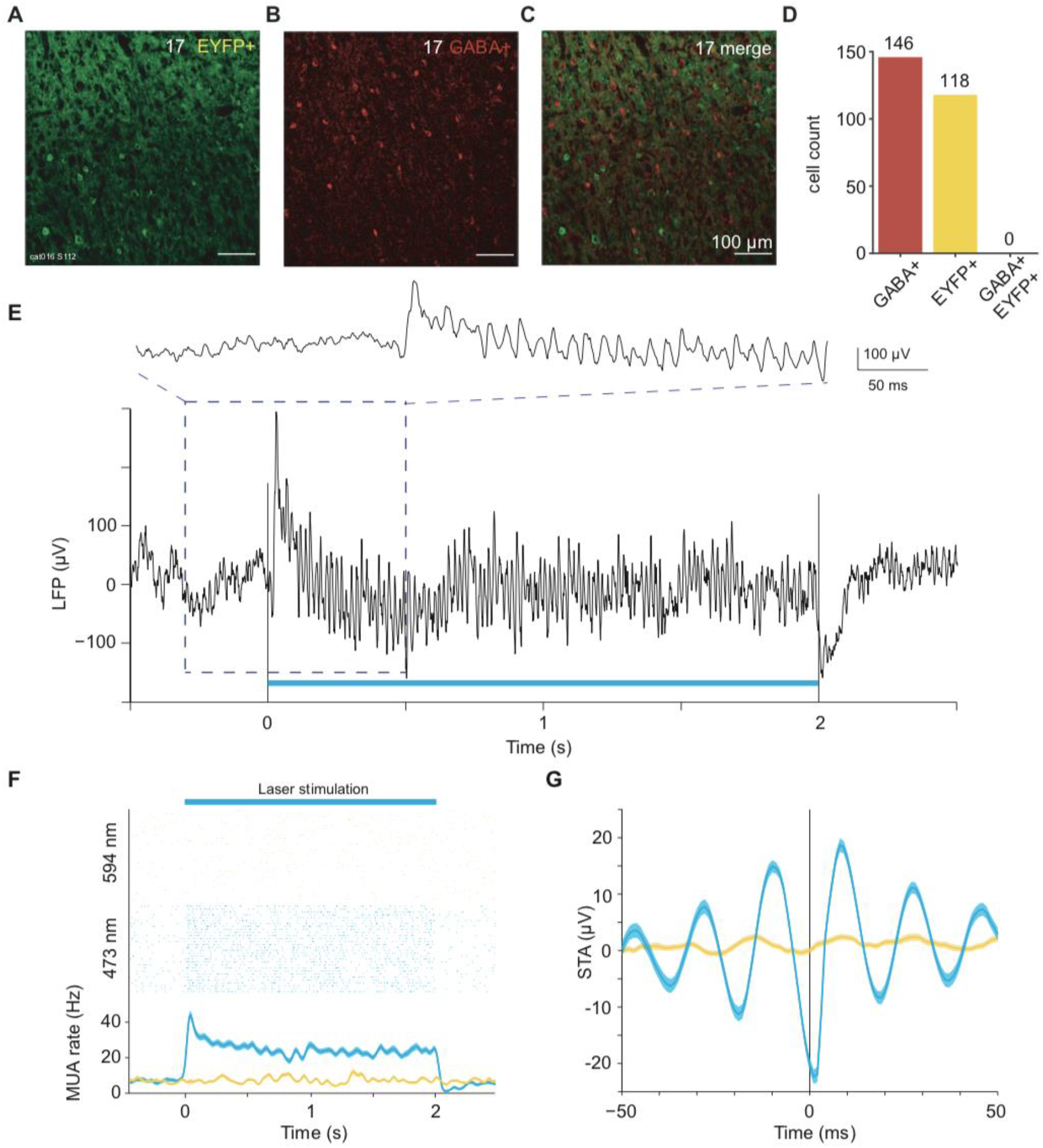
Viral transfection and gamma-band resonance to stimulation. (A-C) Confocal microscopy images of immunohistochemistry performed on slices from area 17 after viral transfection. (A) Endogenous fluorescence of ChR2-eYFP, (B) Fluorescence from secondary antibody after staining for GABA+. (C) Merged images, testing for neuronal co-labeling with ChR2-eYFP and GABA+ antibody. No co-labeled neurons can be found. (D) Counts of GABA+ labeled neurons, EYFP+ labeled neurons, and co-labeled neurons in area 17. (E) Example recording site in area 17 shows strong gamma-band synchronization in the local field potential induced by constant-illumination. (F) Robust MUA response to constant illumination at the same site. Blue: 473 nm wavelength light; Yellow: 594 nm wavelength light. Shaded area indicates ±1 SEM across trials. (G) Spike-triggered LFP for example data shown in E and F. Shaded area indicates ±1 SEM across trials.

We performed terminal experiments under general anesthesia 4-6 weeks after virus injection. The transfected portion of cortex was illuminated with blue or yellow laser light (473 nm or 594 nm), while neuronal spike and local field potential (LFP) activity was recorded. Since ChR2 is a light-activated cation channel, illumination of transfected neurons emulates excitatory synaptic inputs. The external excitatory drive to the network can thus be controlled by modulating the intensity of the illumination. Visual cortex exhibits strong gamma-band synchronization in response to sustained visual stimulation (Gray et al., 1992; Gray and Singer, 1989). Gamma-band synchronization has also been reported during optogenetic activation of excitatory cells in the primary motor cortex of macaque monkeys (Lu et al., 2015), as well as primary somatosensory cortex and hippocampus of the mouse (Adesnik and Scanziani, 2010; Akam et al., 2012; Stark et al., 2014). We have previously observed gamma-band synchronization in response to constant optogenetic stimulation of excitatory neurons in the visual cortex of the anesthetized cat (Ni et al., 2016). We now present a more detailed analysis of this phenomenon. A single trial of the LFP response to optogenetic stimulation with 2 s of constant blue light from area 17 is shown in Figure 1E. The raw LFP trace reveals strong optogenetically induced gamma that emerged immediately after the onset of stimulation. Figure 1F shows the spike responses of this recording site for many interleaved trials of stimulation with blue or yellow light, confirming that activation was selective for blue light. Activation was also specific to regions of cortex expressing ChR2, as laser stimulation with blue or yellow light had no measurable effect for control recordings in non-transfected cortex (Fig. S1I,J). Figure 1G shows the spike-triggered average (STA) of the LFP, demonstrating that optogenetic stimulation induced spikes that were locked to the LFP gamma-band component. Results in area 21a were highly similar and example data are presented in the supplemental information (Fig. S2A-C).

This pattern was found very reliably across recording sites. Stimulation with two seconds of constant blue light, as compared to yellow control light, induced strong enhancements in firing rate, which were sustained for the duration of stimulation (Fig. S2D,G; Wilcoxon rank-sum test = 39581, *p*<0.0001, n =163 sites in 4 cats). The ratio of LFP power during stimulation versus pre-stimulation baseline showed an optogenetically induced gamma-band peak around 70 Hz (Fig. S2E,H; Wilcoxon rank-sum test = 14751, *p*<0.0001, n = 99 sites in 4 cats). Note that the gamma-band peak frequency varied across animals and recording sessions, as shown previously (Ni et al., 2016). The LFP gamma-power changes reflected changes in neuronal synchronization, because optogenetic stimulation also induced strong MUA-LFP locking in the gamma band, as quantified by the MUA-LFP PPC (Fig. S2F,I; Wilcoxon rank-sum test = 9389, *p*<0.0001, n = 84 sites in 4 cats). In addition to the induction of gamma-band synchronization, optogenetic stimulation also reduced LFP power at 4-14 Hz (Fig. S2E; Fig. S2H inset), and MUA-LFP locking at 10-12 Hz (Fig. S2F; Fig. S2I inset). These reductions of lower-frequency synchronization are reminiscent of effects of visual stimulation and/or selective attention in awake macaque area V4 (Fries et al., 2008b; Mitchell et al., 2009).

### Greater excitation increases magnitude and frequency of resonance

We next characterized the bandwidth of the network resonance by varying the excitation in the local network. Models and empirical data have both suggested that the frequency of gamma oscillations can increase with increasing excitation (Jia et al., 2013; Lowet et al., 2017; Ray and Maunsell, 2010; Roberts et al., 2013; Traub et al., 1996). We therefore slowly increased excitation linearly in time (ramp stimulation, 3 seconds) to generate increasing excitation in the local network. A time-frequency plot for an example recording site in area 21a is presented in Figure 2A. We found that the network resonance varied non-linearly with the input excitation. Rather than scaling linearly with light strength, network resonance only began after a critical level of excitation was reached (Fig. 2A and B), as previously established *in vitro* and in models (Börgers et al., 2005; Traub et al., 1996). Power and frequency increased sub-linearly with increasing excitation (Fig. 2B and C). Interestingly, previous studies reported that optogenetic drive of excitatory cells in somatosensory cortex and hippocampus of the mouse with light ramps resulted in gamma-band synchronization with a constant frequency (Adesnik and Scanziani, 2010; Akam et al., 2012), and increasing the slope of the ramp gave rise to higher-frequency synchronization in somatosensory cortex (Adesnik and Scanziani, 2010). To further investigate additional non-linearities in the resonance, we stimulated the network with slow symmetric excitation profiles (single-slow-sine-wave stimulation, 10 seconds). A time-frequency plot for an example recording site in area 21a is presented in Figure 2D. Single-slow-sine-wave stimulation revealed amplitude and frequency hysteresis, with the amplitude and frequency of the network resonance increasing sub-linearly after a critical point of excitation was reached, and slowing down more quickly upon waning excitation (Fig. 2E and F).

**Figure 2.**
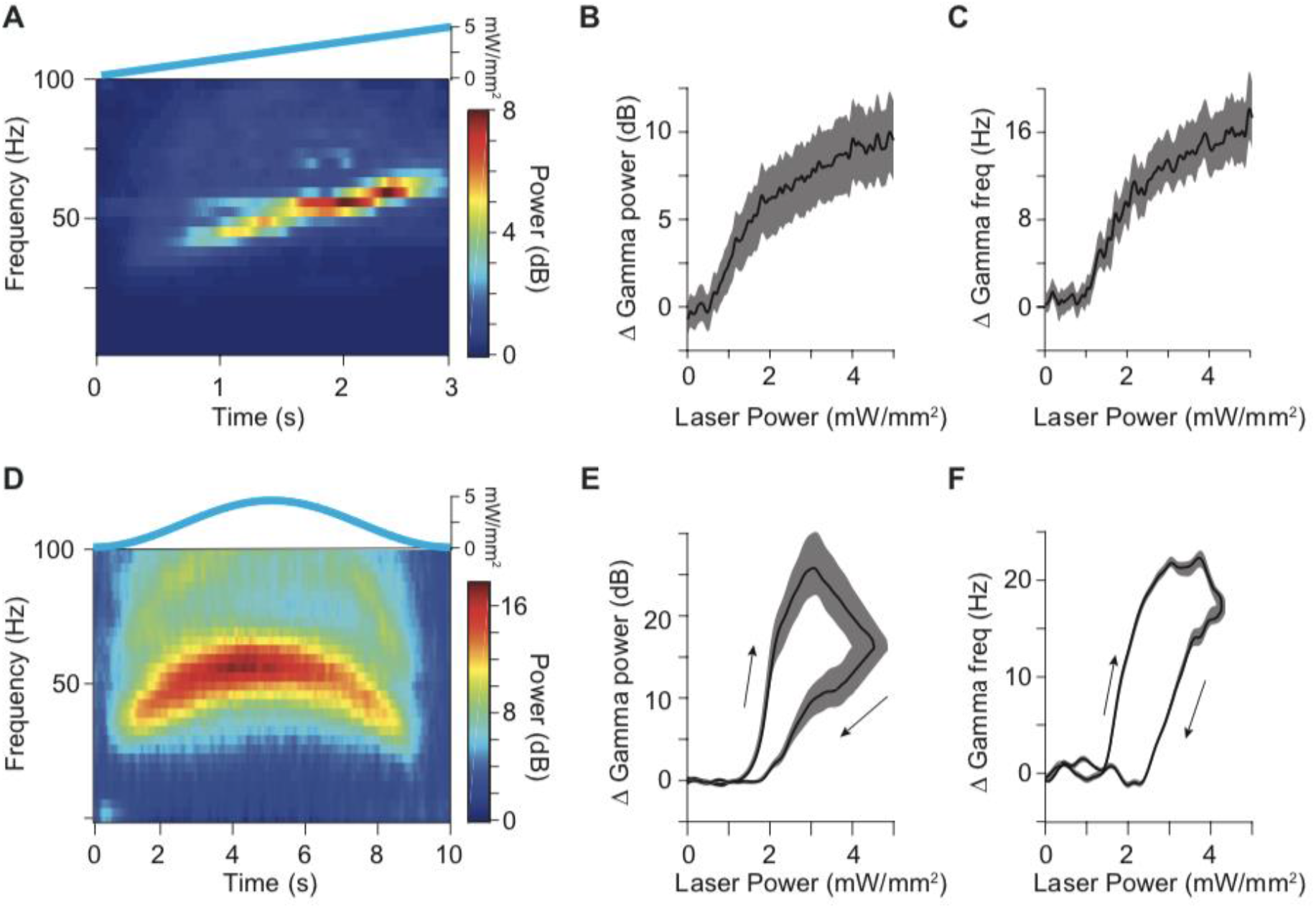
Bandwidth and hysteresis of Gamma-band resonance. (A) Time-frequency plot for an example site in area 21a in response to a slowly increasing ramp stimulus, shown on top. (B) Group result for ramp stimulation shows that the power of the gamma-band resonance increases sublinearly with increasing excitatory drive (N = 58 sites in 5 cats). (C) Same as in B, but for the frequency of the gamma-band resonance. (D) Time-frequency plot for an example site in area 21a to a slow Gaussian temporal profile, shown on top. (E) Group results showing the change in power of gamma-band resonance as a function of laser intensity during slow Gaussian stimulation (N = 52 sites in 5 cats). (F) Same as in E, but for frequency of gamma-band resonance. Arrows indicate hysteresis in response to increasing (upper arrow) versus decreasing (lower arrow) laser power. Shaded areas in panels B, C, E and F indicates ±1 SEM across recording sites.

### Models reveal potential role of non-inactivating M-current in hysteresis

To investigate the network mechanisms underlying the observed resonance phenomena and the hysteresis, we constructed mathematical models of recurrently coupled excitatory and inhibitory neurons. To this end, we used a well-established biophysically realistic pyramidal-interneuron network (PING) model (Börgers, 2017), without additional tuning (Fig. 5A). We initially investigated a model composed of two populations of single-compartment neurons implementing Hodgkin-Huxley dynamics. The excitatory population is based on a simplified model of pyramidal cells (Traub et al., 1991), and the inhibitory population is based on a simplified model of PV+ basket cells (Wang and Buzsáki, 1996). The network has a synaptic model that permits a gradual rise of synaptic gating (Wang, 1999). This model produced strong gamma-band synchronization, as has been reported extensively (Börgers and Kopell, 2003) (Fig. S3A, B).

The PING network reproduced the experimentally observed increase in the power and frequency of the resonance with increased external drive (Fig. S3C,D). Such increases have also been described in vitro (Traub et al., 1996) and in simple networks (Wilson and Cowan, 1972). We implemented a simple leaky-integrate-and-fire (LIF) network, and found that it also exhibited power and frequency increases with increased excitatory drive (Fig. S4A,B). However, neither the PING nor the LIF model were able to reproduce the experimentally observed hysteresis effects (Fig. S3C,D and Fig. S4A,B). We therefore modified the PING model, by adding a non-inactivating M-current to the excitatory population (PING+M model). The M-current is a potassium current that is active at rest and during depolarization and raises the threshold for action potential generation. The PING+M model has lower firing rates and a lower resonant frequency for equal excitatory drive, as compared to the PING model (Fig. S3E,F). The PING+M model was able to produce both power and frequency hysteresis in qualitative concordance with our experimental findings (Fig. S3G,H, as compared to Fig. 2E,F). The hysteresis evident in the PING+M model was considerably less pronounced than what was observed experimentally, suggesting that more factors, such as additional currents, or cell-classes, are likely to contribute to the hysteresis observed *in vivo.*

### Rhythmic input matching resonance is preferentially transmitted

We next returned to the empirical data and sought to investigate whether the output of the local network, assessed by spike output, demonstrates a preference for temporally varying inputs with a timescale matching the network resonance, as has been suggested by computational models (Sherfey et al., 2018). We drove rhythmic excitation in the network with sinusoidal stimulation of 5, 10, 20, 40 and 80 Hz. Light intensity was adjusted per recording site (see Methods) and was kept constant for a given site across the different stimulation frequencies. Sinusoids of all applied frequencies resulted in clear increases in firing rate, with strong rhythmicity at the stimulation frequency (Fig. 3). We calculated spike density functions, subtracted the baseline values and averaged them across recordings sites. Figure 3A shows those average spike densities for 10 Hz, 40 Hz and 80 Hz. Note that 10 Hz stimulation resulted in not only an entrained 10 Hz response, but also bursts of gamma-band synchronization around the peak of excitatory drive, in agreement with a previous report in rodent hippocampus (Butler et al., 2016). Note also that 80 Hz stimulation did not result in simple entrainment to the 80 Hz stimulation, but that the response varies on alternate cycles, exhibiting a prominent sub-harmonic to the driving frequency at 40 Hz that was stable for the entire 2 s stimulation period.

**Figure 3.**
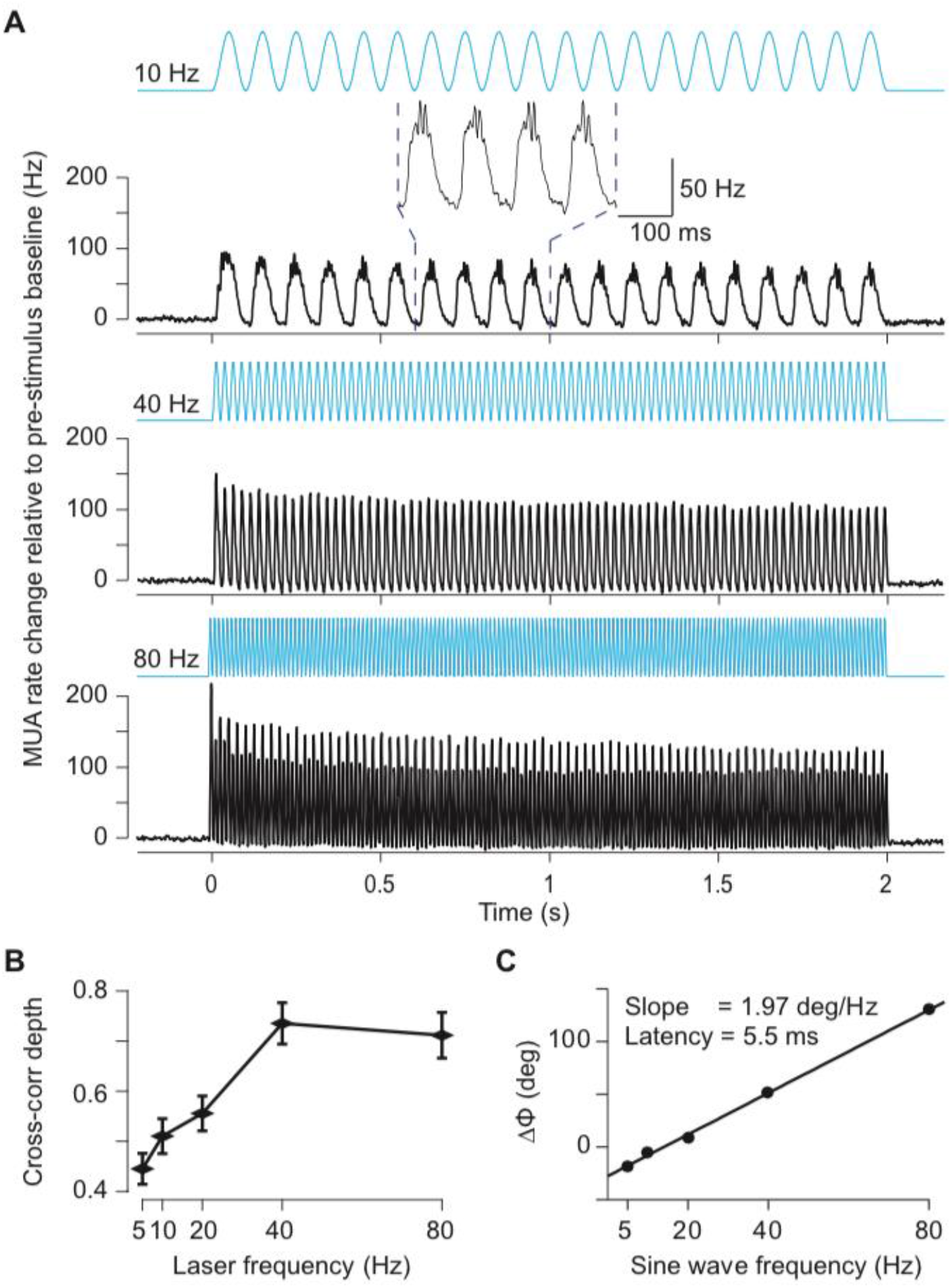
MUA responses to sinusoidal stimulation. (A) MUA spike density (Gaussian smoothing with σ = 1.25 ms and truncated at ±2σ) for 10 Hz (top), 40 Hz (middle) and 80 Hz (bottom) sinusoidal stimulation, respectively. The inset shows an enlarged version of a few cycles to illustrate the gamma-band resonance induced at the peak of the depolarizing phase of the 10 Hz sinusoid. Data were baseline subtracted (−0.5 to 0s) and averaged over all MUA recording sites (N = 60 in 4 cats). Error regions for ±1 SEM across recording sites are smaller than line width. (B) Modulation depth quantified as peak-to-trough distance of the Pearson cross-correlation coefficient as function of the frequency of stimulation. (C) Peak latency from stimulation to MUA response as a function of frequency. The text inset gives the slope and the corresponding latency between optogenetic stimulation and neuronal response.

To capture entrainment by the optogenetic stimulation, we calculated the Pearson cross-correlation coefficient between the respective sinusoid and the resulting spike density, as a function of time lag between the two (Fig. 3B and S5A). We quantified the strength of entrainment as the peak-to-trough distance of the cross-correlation functions (Fig. 3B). Sinusoidal stimulation resulted in entrainment that increased with stimulation frequency to peak at 40 Hz and weakly decreased at 80 Hz (one-way ANOVA, *p* = 1.6E-9, *F*_(4,295)_ = 11.25). The bandwidth of the preferential entrainment matches well the bandwidth found by varying excitation with ramps and gaussian stimulation, and the small fall-off at frequencies above the network resonance is in good agreement with previous modelling work (Sherfey et al., 2018).

Sinusoidal stimulation of different frequencies enabled estimation of neuronal response latencies by computing the slope of relative phases between the stimulation signal and the output MUA across stimulation frequencies (See Supplementary text for an expanded discussion of this method). Figure 3C presents the relative-phase spectrum and reveals a strictly linear relationship, a signature of a fixed time lag. The slope of this linear relationship indicates a latency of 5.5 ms, in good agreement with previous reports (Boyden et al., 2005; Cardin et al., 2009).

### Optogenetic white-noise stimulation reveals causal role of gamma

Finally, and crucially, we emulated input with a white-noise characteristic. White noise realizes continuously unpredictable values (innovation), and thus shows no autocorrelation i.e. no correlation with its own past or future. Thereby, time-lagged correlations between the optogenetically emulated neuronal input and the neuronal spike output cannot be due to time-lagged correlation within the input, but can be unequivocally attributed to a time-lagged correlation between input and output. A time-lagged correlation between an experimentally controlled input and the observed spike output provides direct evidence for a causal role of the input. Importantly, white-noise stimulation enabled us to determine the causal roles separately for each frequency of the spectrum. That is, white-noise excitatory drive during recording of spike output allowed us to determine the directed transfer function of the observed network.

We employed optogenetic stimulation with light intensities following a gaussian random process (sampled at ≈1000 Hz) with a flat power spectrum (Figure 4A, bottom trace). This white-noise stimulus contains the same energy at all frequencies up to 500 Hz. Light intensities were titrated such that firing rates were in the lower half of the dynamic range of the recorded neurons in response to optogenetic stimulation. Figure 4A shows an example LFP and MUA recording for an example trial of white-noise stimulation.

**Figure 4.**
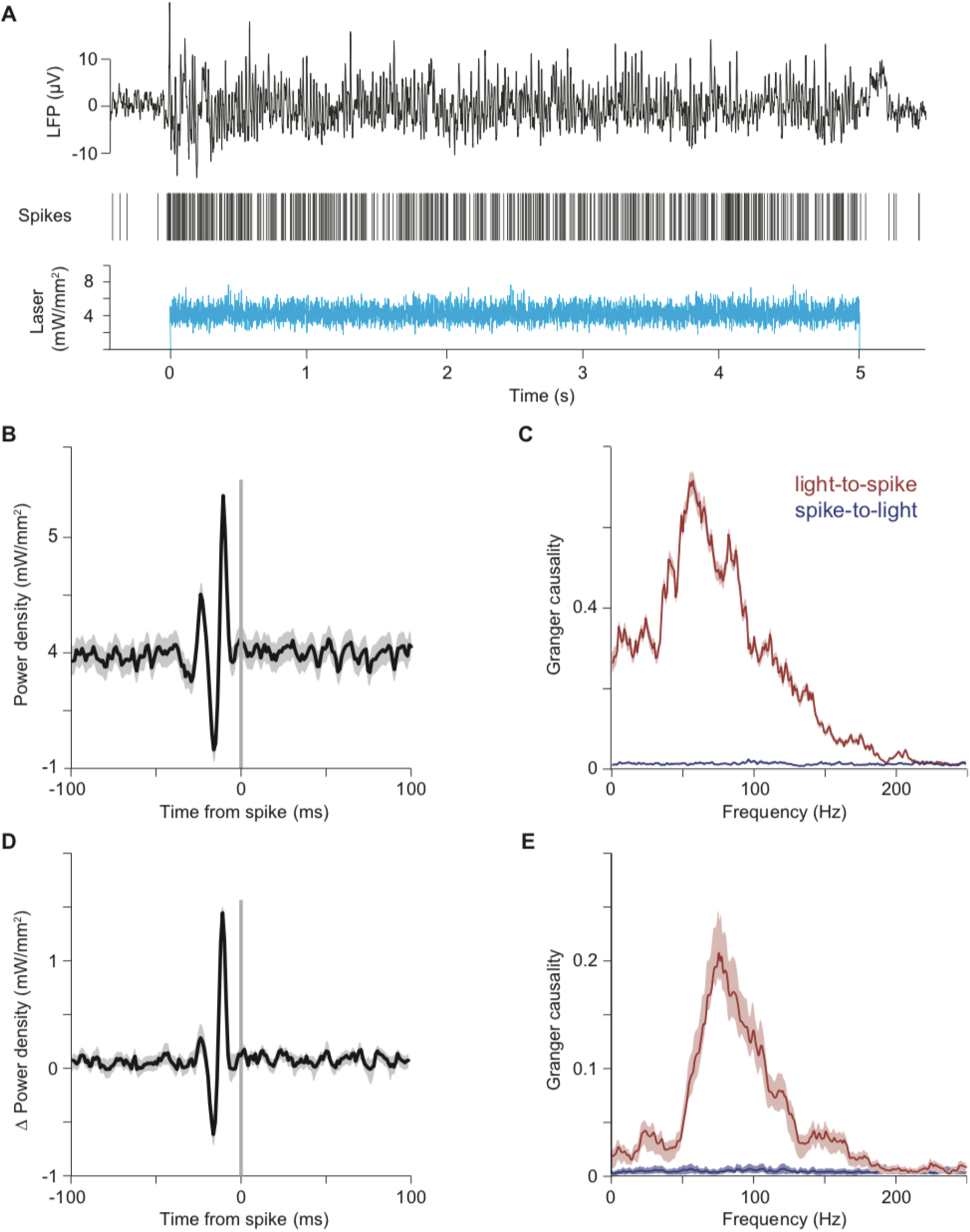
The component of white-noise stimulation coherent with network resonance is transmitted as MUA. (A-C) Example single trial LFP and MUA response to optogenetic white-noise stimulation. The bottom panel shows the white-noise time course of laser intensity. The sequence of vertical lines above it indicates time points of MUA spike occurrence. The black continuous line on top shows the LFP. (B) Spike-triggered average (STA) of laser power density, triggered by the spikes recorded at one example recording site. (C) Granger causality (GC) spectrum for the data shown in (B). Red line shows GC from light to spikes, blue line shows GC from spikes to light (as control). (D, E) Same as (B, C), but for the average across recording sites (N=13 sites in 3 cats).

To reveal the temporal input patterns most reliably driving spikes, we aligned the white-noise time-series that drove the laser to the spikes and averaged it. Figure 4B shows the resulting spike-triggered average light power density for an example recording site. We found that spikes were preceded by a characteristic sequence of increased and decreased light intensity, with a peak-to-peak cycle length corresponding to 75 Hz, suggesting a causal role of the gamma band in eliciting spikes. To quantify this causal influence in a frequency-resolved manner, we calculated the Granger causality of the time-varying light intensity onto the spike train. This revealed a clear peak in the gamma band (Fig. 4C, red). As a control, we also calculated the Granger causality of the spike train onto the light, which confirmed values close to zero, as expected (Fig. 4C, blue). We found very similar effects in the average over recording sites (Fig. 4D,E, N=13 sites in 3 cats), confirming a predominant role of the gamma band in causing spikes.

### Models reveal key role of feedback inhibition in transmission of coherent input

In order to better understand the network behavior under external drive with temporal white noise, we first returned to the PING model without M-current (Fig. 5A). When we stimulated this model with white noise, we found the same signature of frequency dependent transmission as in our experiments (Fig. 5B-C, as compared to Fig. 4B-E). This effect was also evident in the LIF network (Fig. S7A), and in a PING network without I-to-I connectivity (Fig. S7B). The model permitted us to separate excitatory from inhibitory activity, and we found that input in-phase with network excitation, and phase-advanced with respect to network inhibition is preferentially transmitted (Fig. 5B). We computed the Granger causality spectra between the white noise input and the multi-unit activity in the network, and found a high degree of qualitative similarity to our empirical spectra (Fig. 5C, black line; Fig. 4C,E). Again, because we could separate excitation from inhibition in the model, we could separately investigate the transfer from the white-noise input to the excitatory (Fig. 5C, red line) and the inhibitory (Fig. 5C, blue line) activity of the network (see Fig. S7C for corresponding STAs). This suggests that the white-noise components transmitted to network excitation are broader, as compared to the components transmitted to the inhibition. We further investigated the transfer between the excitatory and inhibitory units of the network (Fig. 5D). Excitatory units transmitted variance at gamma, and additionally significant low-frequency variance, to the output of the inhibitory network, whereas inhibitory units transmitted primarily gamma-band components back to the excitatory units (Fig. 5D).

**Figure 5.**
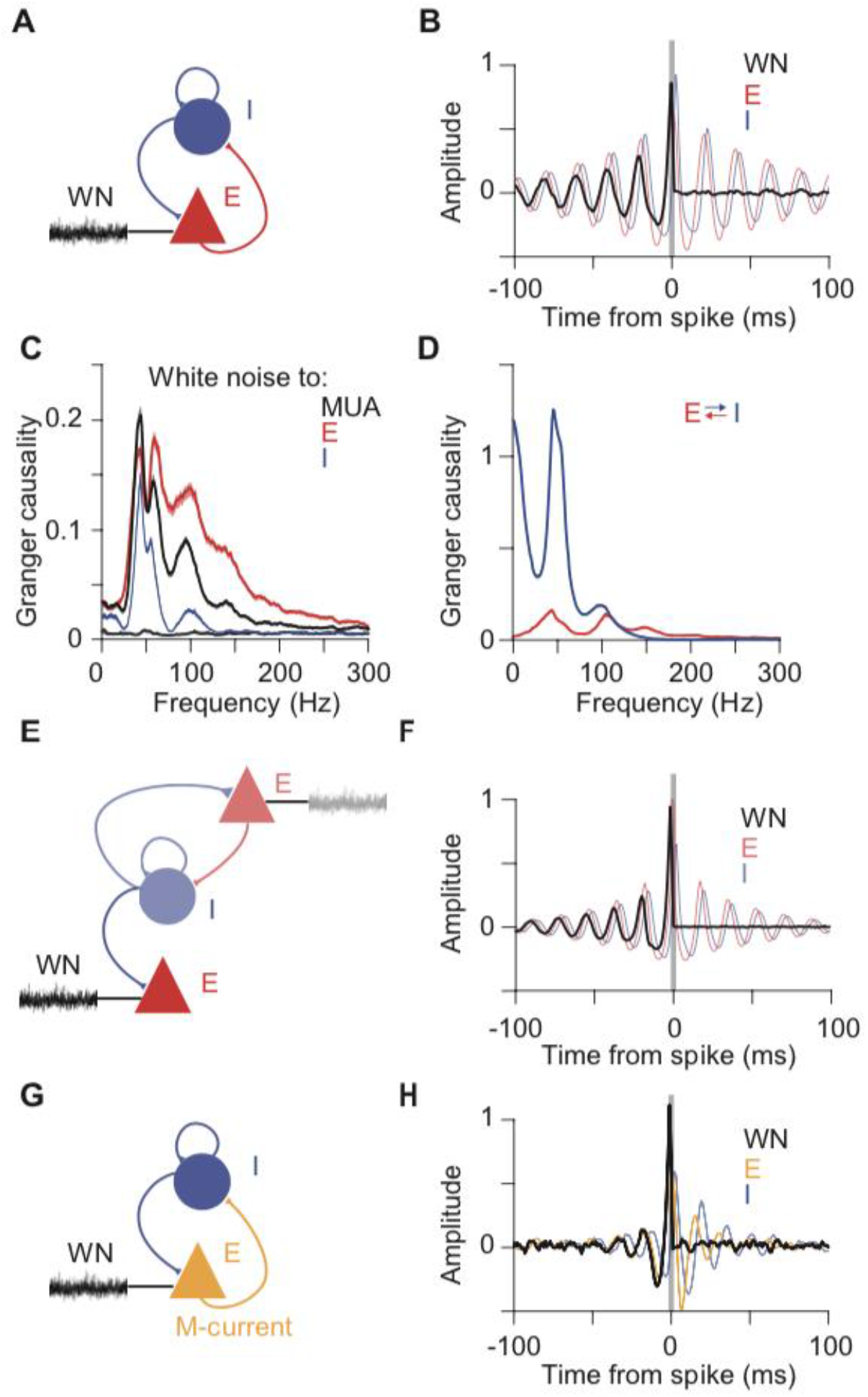
Computational modelling reveals potential mechanism underlying preferential transmission of coherent input. (A) Schematic of PING model driven by white noise. E: Excitatory neuron pool; I: Inhibitory neuron pool; WN: White-noise input. (B) Spike-triggered average of white-noise input signal (black), network excitation (red), and inhibition (blue) demonstrate selective transmission of gamma frequency input matching to the intrinsic dynamics of the network. White-noise averaging was triggered by spikes of all excitatory neurons; results for inhibitory neurons or all neurons (total MUA) are shown in Figure S6E. (C) Granger causality spectrum from white-noise input to total MUA (black), excitatory spikes (red), and inhibitory spikes (blue). Spectra from MUA and spikes to white noise are presented in muted color and overlap near zero. (D) Granger causality spectra between excitation and inhibition in the network. Spectrum from excitatory spikes to inhibitory spikes (blue) and vice-versa (red). (E) Schematic of a two-excitatory-populations model. The PING network shown on top, in lighter colors, contains a first excitatory population and an inhibitory population, and generates gamma upon white-noise input. The resulting rhythmic inhibition is fed into a second excitatory population, shown on the bottom, which is driven by independent white noise. (F) Spike-triggered averages based on spikes from the second excitatory population. Averages display the spike-triggered white noise (black) driving the second excitatory population and illustrate entrainment by the excitation (red) and inhibition (blue) of the recurrently coupled PING network. (G) Schematic of PING+M model driven by white noise. (H) Spike-triggered averages as in panel B, but for the PING+M model.

To further understand the mechanisms of preferential transfer, we next asked whether the excitatory population receiving the white-noise innovation must project to the inhibitory population and thereby entrain a rhythm, or whether a resonant pool, isolated from the white-noise innovation, but projecting inhibitory synapses to that population could implement selective transmission. We simulated a network with one population of inhibitory neurons and two separate populations of excitatory neurons (Fig. 5E). A first excitatory population (illustrated at the top of Fig. 5E) was recurrently connected to the inhibitory population, and when this circuit was driven by white noise input, it generated gamma resonance. The resulting output of the inhibitory population was fed into the second excitatory population (illustrated at the bottom of Fig. 5E), which did not project back to the inhibitory pool. White-noise input to this second excitatory population was preferentially transmitted, if it was coherent with the gamma-rhythmic inhibition (Fig. 5F). Thus, rhythmic gating can be exerted by one circuit onto a separate, gated, circuit.

Finally, because the M-current had been necessary to explain the experimentally observed hysteresis effects, we asked whether selective transmission occurs in the PING+M model (Fig. 5G). Using the same model parameters as used for the investigation of hysteresis, we performed analysis of the network under white-noise stimulation. We found that the PING+M model also exhibited selective transmission (Fig. 5H). Intriguingly, the M-current significantly reduced the timescale of the selective transmission, producing spike-triggered white noise in better agreement with that found experimentally (Fig. 5H as compared to Fig. 4B,D). Together with the hysteresis results, the close qualitative match between the experimental and the PING+M spike-triggered white-noise results suggests the influence of some form of spike-frequency adaptation. As mentioned above, it is likely that additional cell-classes, or conductances may play a role *in vivo* and require further investigation. In any case, given the potential role of the M-Current suggested by our findings, it would be interesting to investigate the impact of acetylcholine on the phenomena described here. Acetylcholine can have an antagonistic effect on the M-current via muscarinic receptors, and in the PING+M model this would increase the power and frequency of the circuit resonance, enhance the amplitude and timescale of selective transmission, and reduce the hysteresis of the gamma-band resonance. Intriguingly, a number of previous studies in cat visual cortex have already described increased gamma band synchronization after electrical stimulation of the midbrain reticular formation (Munk et al., 1996), which appears to depend on muscarinic receptors (Rodriguez et al., 2004).

## Discussion

Visual stimulation induces clear gamma-band synchronization in cat visual cortex, both during wakefulness (Fries et al., 2002; Gray and Viana Di Prisco, 1997) and anesthesia (Gray et al., 1992). We recorded LFPs and neuronal spike output in the visual cortex of anesthetized cats, while optogenetically emulating external, excitatory inputs to pyramidal neurons with precise experimental control. Controlling external excitatory drive allowed us to investigate the functional consequences of the cortical gamma-band resonance. Optogenetic excitation with a variety of temporal patterns produced gamma-band activity qualitatively similar to that found for visual stimulation. A better understanding of cortical resonance sheds light on the dynamic transformations performed by the local circuit and reveals how time-varying excitation is transmitted.

We confirmed that visual cortex transforms constant excitation into strong gamma-band synchronization, producing rhythmic spike output similar to visual stimulation (Ni et al., 2016). Slowly increasing excitation with ramps increased the strength and frequency of synchronization, and revealed a threshold of excitation necessary for the ignition of synchronization. A positive correlation between excitatory drive and the strength and frequency of gamma-band synchronization has been predicted by computational models, demonstrated *in vitro*, and is reminiscent of effects seen *in vivo* for visual contrast and salience (Fries, 2015; Hadjipapas et al., 2015; Jia et al., 2013; Lowet et al., 2017; Ray and Maunsell, 2010; Roberts et al., 2013; Traub et al., 1996). Slow, temporally symmetric excitation profiles demonstrated profound hysteresis in both the strength and frequency of the synchronization. While hysteresis in synchronization has so far been unreported to our knowledge, it is reminiscent of effects seen when visual contrast is symmetrically varied (for example, see Fig. 3 of (Ray and Maunsell, 2010)). Modelling indicated that hysteresis could arise from spike-frequency adaptation via a non-inactivating potassium current (M-current), suggesting that acetylcholine effects on the M-current may modify the dynamics of gamma band resonance (Börgers et al., 2005; Fellous and Sejnowski, 2000; Fisahn et al., 1998; Munk et al., 1996; Rodriguez et al., 2004). The observed hysteresis could play a powerful role in differentiating populations of cells with increasing versus decreasing excitation, even if the total level of excitation in the populations is equal. Future studies should elucidate the rich, non-linear features of the resonance described here, such as the minimal excitatory drive required for resonance, its dynamic range, and its interaction with neuromodulatory signals.

Varying external drive on faster timescales enabled us to investigate how cortical resonance selectively transmits components of dynamic input. The effect of the network resonance on variable input was first demonstrated for rhythmic, sinusoidal excitation. Sinusoidal drive was transformed by the network into spike output with a fidelity that increased up to 40 Hz and declined slightly for 80 Hz. Intriguingly, slow sinusoidal input gave rise to bursts of gamma band synchronization at the peaks.

Crucially, the precise temporal control afforded by optogenetics enabled characterization of the network response to stochastic, white noise sequences. White-noise stimulation facilitated causal analysis of network transmission: from external excitatory input to neuronal spike output. The gamma-band component of the stochastic input preferentially drove spiking in the neuronal population. Thus, feline visual cortex is predisposed to transform external excitation with a variety of temporal profiles into gamma-rhythmic spike output. Further, the resulting gamma-rhythmic output is ideally suited to preferentially drive activity in downstream populations.

Network resonance emerges from the interaction between excitatory and inhibitory elements. In computational models, including those presented here, network resonance is determined largely by feedback inhibition. While resonance arises in reduced models with homogeneous cellular properties, the cat visual cortex contains a great deal of heterogeneity. The dominant gamma-band resonance we observed could be due to intracellular mechanisms, network properties, or combinations of both. Intracellular transfer functions have been characterized for assorted cell classes using *in vitro* electrophysiology and optogenetics. While there is diversity depending on morphology and channel composition, the dominant cell class we drove with light, pyramidal cells, typically exhibits a low-pass characteristic. Previous work characterized the transfer function of a variety of opsins, including the opsin used here (hChR2(H134R)), in cultured pyramidal cells and found that transfer peaked at 3 Hz and declined smoothly for higher frequencies, with currents reduced by half at ~40 Hz (ChR2R in Fig. 1 of (Tchumatchenko et al., 2013)). Therefore, the gamma-band resonance observed in the present study is most likely not due to the opsin or electrical properties of the individual neurons, but rather predominantly determined by feedback inhibition in the network (Buzsáki and Wang, 2012). This network mechanism is likely assisted and amplified by cellular mechanisms. Interneurons can show 1:1 phase locking to suprathreshold sinusoidal current injections up to 50 Hz (Fellous et al., 2001). When the transfer function from injected current to spike times is directly measured for cortical interneurons in slices of ferret prefrontal cortex, it reveals a broad peak in the gamma range (Hasenstaub et al., 2005). Additionally, specialized classes of excitatory neurons have been described in cat and macaque visual cortex, with properties that likely promote gamma-band resonance (Gray and McCormick, 1996; Onorato et al., 2020).

Interestingly, the spike-triggered average revealed that spikes were preceded not only by rhythmic peaks, but also by rhythmic troughs, suggesting that input which matches the intrinsic timescale of feedback inhibition is preferentially transmitted. In a driven state, network excitation and inhibition wax and wane with a delay determined by features of synaptic connectivity. This creates windows of enhanced susceptibility to external drive, and the pace of network inhibition will preferentially permit excitatory cells to transmit components of their time-varying extrinsic drive that match the endogenous dynamics (Fries, 2015). Exogenously driven excitatory spikes will subsequently drive inhibitory neurons and renew the cycle of feedback inhibition. If excitation arrives out of phase with the network rhythm, it can prematurely drive inhibition in a feedforward manner, and sufficient premature capture of inhibition will lead to desynchronization of the inhibitory pool. Such premature forcing is kept in check by the strong synchronization within the inhibitory pool, via dense I-I coupling. Thus, exogenous excitation competes with the endogenous pace set by strong feedback inhibition.

Spike-triggered average (STA) analysis has been used to characterize the input-output relationship of single neurons, both in terms of their receptive field properties (Chichilnisky, 2001; Pillow et al., 2008), and in terms of their resonance properties (Bryant and Segundo, 1976; Mainen and Sejnowski, 1995; Marmarelis and Naka, 1972). It is also routinely used to estimate the locking of neurons to simultaneous population activity, either by spike-triggered LFP averaging (Fries et al., 1997), or spike-triggered covariance analysis (Pillow et al., 2008). STA analysis of both intracellularly recorded membrane potentials (Azouz and Gray, 2008; Hasenstaub et al., 2005) and LFPs (Fries et al., 1997), has revealed strong gamma-band phase-locking during visual stimulation. As membrane potentials and LFPs reflect synaptic currents (Pesaran et al., 2018), these observations are consistent with a scenario in which spikes are specifically caused by the gamma component of synaptic inputs. However, these findings are also consistent with a scenario in which visual stimulation induced gamma-rhythmic neuronal activity reflected in both spiking and LFP, without a specific causal role of gamma-rhythmic inputs. Optogenetic white noise stimulation allowed us to isolate the effect of external gamma-rhythmic drive from ongoing synchronization. We were therefore able to demonstrate the causal role of network resonance in selectively transmitting the gamma component of time-varying external input. Importantly, the gamma-rhythmic component of the spike-triggered white-noise average cannot be explained by the mere fact that the stimulation induced gamma-rhythmic neuronal spiking. Rather, it required that spikes were time locked (and thereby phase locked) to the relevant temporal pattern in the white noise. If white noise had simply induced spikes that were gamma-rhythmic but not phase-locked to the gamma component of the white noise, the STA of the white noise would have been flat. However, the STA revealed significant modulation in the gamma band, suggesting that spikes were preferentially driven by the input’s gamma components.

The gamma synchronization produced by white-noise input was weaker, and more unstable than that produced with constant stimulation (Fig. S6). During constant stimulation, the exogenous drive lacks temporal structure, and network dynamics are dominated by the endogenous resonance. However, during white-noise stimulation, endogenous dynamics are perturbed by broadband exogenous drive, resulting in irregular, fragmented synchronization. Similarly, gamma-band activity in macaque V1 is strong when induced by a smoothly moving grating, and substantially reduced by the addition of random motion (Kruse and Eckhorn, 1996). Interestingly, temporally variable exogenous drive leads to precise spike timing, increased stimulus information, and improved perceptual discrimination (Buracas et al., 1998; Christensen et al., 2019; Mainen and Sejnowski, 1995). Complementary results suggest that endogenous gamma dynamics provide additional temporal structure that can enhance the information communicated by neurons (Azouz and Gray, 2003; Harris et al., 2003; Womelsdorf et al., 2012). Together, these results suggest that networks balance the deviations introduced by exogenous drive with the timescale imposed by their endogenous dynamics. Indeed, exogenous transients may function as an external clock to synchronize activity and facilitate transmission, while under continuous or slowly varying drive, resonance may assume the role of timekeeper and discretize transmission into synchronous packages so as to maximize their effect on downstream populations. Under such a regime, temporal information imposed by a variable stimulus will be faithfully conveyed, and in the absence of exogenous temporal structure, the synchronization imposed by network resonance will endow neuronal communication with increased reliability and precision (Fries, 2015). The balance of exogenous and endogenous drive is likely to fluctuate dynamically according to their relative strength, or other variables which can alter the dynamic set-point of the circuit. The flexible balancing of extrinsic and intrinsic factors provides a powerful means to selectively amplify and propagate or suppress and gate sensory signals according to behavioral state or goals.

The experiments reported here were limited to visual cortex and have focused on the gamma-band resonance prominent in activated visual cortex (Brunet et al., 2015; Gray and Singer, 1989; Onorato et al., 2020). However, all recurrently coupled excitatory-inhibitory networks are likely to demonstrate similar resonances, which will function to selectively filter their input and temporally tune their output. This reasoning predicts that spikes in other areas, in which other rhythms predominate (Brown et al., 1998; Csicsvari et al., 2003; Fries, 2009; Gregoriou et al., 2009; Pesaran et al., 2002), might be caused predominantly by the corresponding rhythm in their input. Likewise, because our experiments were carried out in anesthetized animals, we could not establish the behavioral relevance of the reported phenomena. Previous work has used white noise flicker to investigate the reverberatory nature of visual responses (VanRullen and Macdonald, 2012) and attentional gating of stimulus information (Grothe et al., 2018). These promising results suggest that optogenetic stimulation in behaviorally engaged circuits may provide a powerful means to probe the dynamic routing of information between relevant brain areas.

The filtering and preferential transmission reported here suggest that resonance is a compelling mechanism by which to achieve flexible communication. The resonant frequency of a circuit or population will determine the communication channel of that circuit, and coherent input will be transmitted, while non-coherent input is suppressed (Akam and Kullmann, 2010). Indeed, distinct resonances are likely to exist within a single cortical area, for example, between distinct neuronal subpopulations, projections, or laminae. For example, superficial and deep layers in macaque areas V1, V2 and V4 show very different rhythms during activation. While superficial layers express strong gamma synchronization, deep layers show an alpha-beta rhythm (Buffalo et al., 2011; van Kerkoerle et al., 2014). Rhythms can also change dynamically depending on intrinsic or extrinsic factors such as behavioral state or cognitive context, and such changes might alter resonances and input-output functions, perhaps via modulatory signals (Gulbinaite et al., 2019). A hierarchy of areas with intrinsic resonances could act to selectively distinguish and propagate feedforward and feedback signals in the spectral domain, as has been suggested by functional-anatomical studies (Bastos et al., 2015; Michalareas et al., 2016; van Kerkoerle et al., 2014) and modelling (Lee et al., 2013). It will be a highly interesting task for future studies to probe resonances in different areas, layers, projections, or cell classes and especially in different cognitive contexts. Note that the approach presented here can also be used to investigate the transfer between input to one neuronal group and the spike output of another neuronal group, with the two groups possibly residing in different layers and/or areas. With recordings at site A and stimulation at sites B and C, it might be possible to characterize not only the spectral transfer function from B to A, but also the frequency-resolved modulatory influence of C on this transfer function. By facilitating such investigations, the presented approach provides a novel framework in which to study the mechanisms underlying flexible neuronal communication.

## Acknowledgements

We thank Gustavo Rohenkohl for his assistance and helpful suggestions during a portion of the experiments reported here. This work was supported by DFG (SPP 1665 FR2557/1-1, FOR 1847 FR2557/2-1, FR2557/5-1-CORNET, FR2557/6-1-NeuroTMR, FR2557/7-1-DualStreams to P.F.; EXC 1086, DI 1908/5-1, DI 1908/6-1 to I.D.), BMBF (01GQ1301 to I.D.), EU (HEALTH-F2-2008-200728-BrainSynch, FP7-604102-HBP, FP7-600730-Magnetrodes to P.F.; ERC Starting Grant OptoMotorPath to I.D.), a European Young Investigator Award to P.F., the FENS-Kavli Network of Excellence to I.D., National Institutes of Health (1U54MH091657-WU-Minn-Consortium-HCP to P.F.), the LOEWE program (NeFF to P.F. and I.D.).

## Author contributions

C.M.L., J.N, T.W., P.F. designed research; C.M.L., J.N, T.W., P.J., I.D., P.F. performed experiments; C.M.L., J.N., T.W., P.F. analyzed data; C.M.L. and A.L. performed modelling with input from P.F.; C.M.L., J.N., P.F. wrote the paper.

## Methods

Eight adult domestic cats (*felis catus*; four females) were used in this study. We used cats because the physiology with regard to gamma is highly similar to human and non-human primates (Fries et al., 2008a), both during wakefulness (Fries et al., 2002; Gray and Viana Di Prisco, 1997) and light anesthesia (Gray et al., 1992). Data from the same animals were used in a previous study (Ni et al., 2016). All procedures complied with the German law for the protection of animals and were approved by the regional authority (Regierungspräsidium Darmstadt). After an initial surgery for the injection of viral vectors and a 4-6 week period for opsin expression, recordings were obtained during a terminal experiment under general anesthesia.

### Viral vector injection

For the injection surgery, anesthesia was induced by intramuscular injection of ketamine (10 mg/kg) and dexmedetomidine (0.02 mg/kg), cats were intubated, and anesthesia was maintained with N_2_O:O_2_ (60/40%), isoflurane (~1.5%) and remifentanil (0.3 μg/kg/min). Four cats were injected in area 17 and another four cats in area 21a. Rectangular craniotomies were made over the respective areas (Area 17: AP [0, −7.5] mm; ML: [0, 5] mm; area 21a: AP [0,-8] mm, ML [9, 15] mm). The areas were identified by the pattern of sulci and gyri, and the dura mater was removed over part of the respective areas. Three to four injection sites were chosen, avoiding blood vessels, with horizontal distances between injection sites of at least 1 mm. At each site, a Hamilton syringe (34 G needle size; World Precision Instruments) was inserted with the use of a micromanipulator and under visual inspection to a cortical depth of 1 mm below the pia mater. Subsequently, 2 μl of viral vector dispersion was injected at a rate of 150 nl/min. After each injection, the needle was left in place for 10 min before withdrawal, to avoid reflux. Upon completion of injections, the dura opening was covered with silicone foil and a thin layer of silicone gel, the trepanation was filled with dental acrylic, and the scalp was sutured.

We first tried to transfect with AAV5, because this serotype had been successfully used in many studies on different species (Diester et al., 2011). In one cat, area 17 of the left hemisphere was injected with AAV5-CamKIIα-ChR2-eYFP (titer 4*10^13^ GC/ml). However, this did not result in detectable ChR2-eYFP expression. This failure of AAV5 expression is consistent with one previous study suggesting that AAV5 is not able to provide transduction in the cerebral cortex of the cat (Vite et al., 2003). Subsequently, we tried both AAV1 and AAV9 and found robust transfection with both of these serotypes. In one cat, area 17 in the left hemisphere was injected with AAV1-CamKIIα-hChR2(H134R)-eYFP (titer 8.97*10^12^ GC/ml) and area 17 in the right hemisphere with AAV9-CamKIIα-ChR2-eYFP (titer 1.06*10^13^ GC/ml). In two cats, area 17 of the left hemisphere was injected with AAV1-CamKIIα-hChR2(H134R)-eYFP (titer: 1.22*10^13^ GC/ml). In four cats, area 21a of the left hemisphere was injected with AAV9-CamKIIα-hChR2(H134R)-eYFP (titer: 1.06*10^13^ GC/ml). The DNA plasmids were provided by Dr. Karl Deisseroth (Stanford University, Stanford, CA). AAV5 viral vectors were obtained from UNC Vector Core (UNC School of Medicine, University of North Carolina, USA); AAV1 and AAV9 viral vectors were obtained from Penn Vector Core (Perelman School of Medicine, University of Pennsylvania, USA).

### Neurophysiological recordings

For the recording experiment, anesthesia was induced and initially maintained as during the injection surgery, only replacing intubation with tracheotomy and remifentanyl with sufentanil. After surgery, during recordings, isoflurane concentration was lowered to 0.6%-1.0%, eye lid closure reflex was tested to verify narcosis, and vecuronium (0.25mg/kg/h i.v.) was added for paralysis during recordings. Throughout surgery and recordings, Ringer solution plus 10% glucose was given (20 ml/h during surgery; 7 ml/h during recordings), and vital parameters were monitored (ECG, body temperature, expiratory gases).

Each recording experiment consisted of multiple sessions. For each session, we inserted either single or multiple tungsten microelectrodes (~1 MΩ at 1 kHz; FHC), or three to four 32-contact probes (100 μm inter-contact spacing, ~1 MΩ at 1 kHz; NeuroNexus or ATLAS Neuroengineering). In one cat, one 16-contact probe with 150 μm inter-contact spacing and one 46 μm optic fiber, and one 16-contact probe with 150 μm inter-contact spacing and four 46 μm optic fibers were used (Plexon V- and U-probe, respectively). Standard electrophysiological techniques (Tucker Davis Technologies, TDT) were used to obtain multi-unit activity (MUA) and LFP recordings. For MUA recordings, the signals were filtered with a passband of 700 to 7000 Hz, and a threshold was set to retain the spike times of small clusters of units. For LFP recordings, the signals were filtered with a passband of 0.7 to 250 Hz and digitized at 1017.1 Hz.

### Photo-stimulation

Optogenetic stimulation was done with a 473 nm (blue) laser or with a 470 nm (blue) LED (Omicron Laserage). A 594 nm (yellow) laser was used as control. Laser light was delivered to cortex through a 100 μm or a 200 μm diameter multimode fiber (Thorlabs), LED light through a 2 mm diameter polymer optical fiber (Omicron Laserage). Fiber endings were placed just above the cortical surface, immediately next to the recording sites with a slight angle relative to the electrodes. Laser waveform generation used custom circuits in TDT, and timing control used Psychtoolbox-3, a toolbox in MATLAB (MathWorks) (Brainard, 1997).

For white noise stimulation, the laser was driven by normally distributed white noise, with light intensities updated at a frequency of 1017.1 Hz. For each recording session, the mean of the normal distribution was chosen to fall into the lower half of the dynamic range of the laser-response curve of the recorded MUA. This resulted in mean values in the range of 3-12 mW/mm2 (13 MUA recording sites in the 3 cats showing expression of ChR2 in area 17). The standard deviation (SD) of the normal distribution was scaled to be 1/2 the mean. The resulting distributions were truncated at 3.5 SDs. The resulting range of laser intensities always excluded both zero and maximal available laser intensities and thereby avoided clipping.

### Histology

After conclusion of recordings, approximately five days after the start of the terminal experiment and still under narcosis, the animal was euthanized with pentobarbital sodium and transcardially perfused with phosphate buffered saline (PBS) followed by 4% paraformaldehyde. The brain was removed, post-fixed in 4% paraformaldehyde and subsequently soaked in 10%, 20% and 30% sucrose-PBS solution, respectively, until the tissue sank. The cortex was sectioned in 50 μm thick slices, which were mounted on glass slides in antifade medium, protected with coverslips, and subsequently imaged with a confocal laser scanning microscope (CLSM, Nikon C2 90i, Nikon Instruments) for eYFP-labelled neurons.

#### Immunohistochemistry

In two cats, one with injections in area 17 and one with injections in area 21a, slices were processed as described above and additionally stained for parvalbumin (PV) and gamma-Aminobutyric acid (GABA). To this end, slices were preincubated in 10% normal goat serum (NGS) with 1% bovine serum albumin (BSA) and 0.5% Triton X-100 in phosphate buffer (PB) for 1 h at room temperature to block unspecific binding sites. Floating slices were stained for PV (overnight, rabbit anti-Parvalbumin, NB 120-11427, Novus Biologicals) and GABA (48 hours, rabbit anti-GABA, ABN131, Merck Millipore) in 3% NGS containing 1% BSA and 0.5% Triton X-100. After washing two times 15 min in PB, the slices were incubated with the secondary antibody (goat anti-rabbit Alexa Fluor 647, A-21244, Thermo Fisher Scientific) in 3% NGS containing 1% BSA and 0.5% Triton X-100 for 1 h at room temperature. Finally, slices were again washed in PB, protected with coverslips and imaged with a Zeiss CLSM, using a 25X water immersion objective.

### Data analysis

All data analysis was performed using custom code and the Fieldtrip toolbox (Oostenveld et al., 2011), both written in MATLAB (MathWorks).

Spike densities, MUA-laser cross-correlation, LFP power spectra, and MUA-LFP PPCs. MUA rate was smoothed with a Gaussian (for constant light stimulation: SD = 12.5 ms; for stimulation with pulse trains and sinusoids: SD = 1.25 ms; in each case truncated at ± 2 SD) to obtain the spike density.

To quantify the locking of neuronal responses to optogenetic stimulation, we calculated the Pearson correlation coefficient between MUA spike density and laser intensity as a function of time shift between them.

LFP power spectra were calculated for data epochs that were adjusted for each frequency to have a length of 4 cycles and moved over the data in a sliding-window fashion in 1 ms steps. Each epoch was multiplied with a Hann taper, Fourier transformed, squared and divided by the window length to obtain power density per frequency. For the different stimulation frequencies f, LFP power is shown as ratio of power during stimulation versus pre-stimulation baseline (−0.5 s to −0.2 s relative to stimulation onset).

MUA-LFP locking was quantified by calculating the MUA-LFP PPC (pairwise phase consistency), a metric that is not biased by trial number, spike count or spike rate (Vinck et al., 2010). Spike and LFP recordings were always taken from different electrodes. For each spike, the surrounding LFP was Hann tapered and Fourier transformed. Per spike and frequency, this gave the MUA-LFP phase, which should be similar across spikes, if they are locked to the LFP. This phase similarity is quantified by the PPC as the average phase difference across all possible pairs of spikes. To analyze PPC as a function of frequency and time (Fig. 4 and 9), the LFP around each spike in a window of ±2 cycles per frequency was Hann tapered and Fourier transformed. PPC was then calculated for epochs of 100 ms length, i.e. using the phases of spikes in those epochs, moved over the data in a sliding-window fashion in 1 ms steps. To analyze PPC with higher spectral resolution (Fig. 5), the LFP around each spike in a window of ±0.5 s (Fig. 5F, lower frequencies) or ±0.25 s (Fig. 5F, higher frequencies) was Hann tapered and Fourier transformed to obtain the spike phase. For a given MUA channel, MUA-LFP PPC was calculated relative to all LFPs from different electrodes and then averaged.

#### Estimation of Granger causality (GC) between light time course and MUA spike trains

The GC spectrum was first estimated separately for each recording site and subsequently averaged over sites. For each trial, we estimated the Fourier transforms of the input (laser) and the output (MUA). Specifically, each trial was segmented into non-overlapping epochs of 500 ms length. Per epoch, the time series of the input and the output were multiplied with a Hann taper, they were zero-padded to a length of 1000 ms, and their Fourier transforms (FTs) were obtained. The FTs were used to calculate the power-spectral densities (PSDs) of the input and of the output, and the cross-spectral density (CSD) between input and output. CSDs and PSDs were averaged over trials and used for the estimation of GC by means of non-parametric spectral matrix factorization (Dhamala et al., 2008). For the example GC spectrum (Fig. 6C), the error region was determined by a bootstrap procedure, with 100 iterations, each time randomly choosing 30% of the trials. The shown error boundary is the region containing 95% of the bootstrapped estimates. For the average GC spectrum (Fig. 6E), the error region indicates the standard error of the mean across the recording sites.

#### Statistical testing

All inferences were based on the combined data of all animals, for which a given experiment was performed. The resulting inferences are limited to the studied sample of animals, as in most neurophysiological in-vivo studies.

High-resolution spectra of LFP power changes and MUA-LFP PPC were compared between stimulation with blue light and control stimulation with yellow light (Fig. 3B,C). We calculated paired t-tests between spectra obtained with blue and yellow light, across recording sites. Statistical inference was not based directly on the t-tests (and therefore corresponding assumptions will not limit our inference), but the resulting t-values were merely used as a well-normalized difference metric for the subsequent cluster-based non-parametric permutation test. For each of 10,000 permutations, we did the following: 1) We made a random decision per recording site to either exchange the spectrum obtained with blue light and the spectrum obtained with yellow light or not; 2) We performed the t-test; 3) Clusters of adjacent frequencies with significant t-values (p<0.05) were detected, and t-values were summed over all frequencies in the cluster to form the cluster-level test statistic. 4) The maximum and the minimum cluster-level statistic were placed into maximum and minimum randomization distributions, respectively. For the observed data, clusters were derived as for the randomized data. Observed clusters were considered significant if they fell below the 2.5th percentile of the minimum randomization distribution or above the 97.5th percentile of the maximum randomization distribution (Maris and Oostenveld, 2007). This corresponds to a two-sided test with correction for the multiple comparisons performed across frequencies (Nichols and Holmes, 2002).

### PING model

The neurons in the PING model are Hodgkin-Huxley-like point neurons. The excitatory population consists of a simplified version of model pyramidal neurons introduced by(Traub et al., 1991), the reduced Traub-Miles (RTM). The inhibitory population consists of model basket cells introduced by (Wang and Buzsáki, 1996). The parameters for the model are presented in the tables below, and we refer to the original publication of the model for more details (Börgers, 2017).

PING Neuron parameters:

| | C ($\mu\text{F}/\text{cm}^2$) | V<sub>Na</sub> (mV) | V<sub>K</sub> (mV) | V<sub>L</sub> (mV) | g<sub>Na</sub> (mS/cm<sup>2</sup>) | g<sub>K</sub> (mS/cm<sup>2</sup>) | g<sub>L</sub> (mS/cm<sup>2</sup>) |
| --- | --- | --- | --- | --- | --- | --- | --- |
| E (RTM) | 1 | 50 | -100 | -67 | 100 | 80 | 0.1 |
| I (WB) | 1 | 55 | -90 | -65 | 35 | 9 | 0.1 |

PING Network parameters:

|  |  |
| --- | --- |
| N<sub>E</sub> | 200 |
| $N_I$ | 50 |
| $I_E$ | $1.5 \mu\text{A}/\text{cm}^2$ |
| $\sigma_E$ | $0.05 \mu\text{A}/\text{cm}^2$ |
| $I_I$ | $0 \mu\text{A}/\text{cm}^2$ |
| $g_{EE}$ | $0 \text{ mS}/\text{cm}^2$ |
| $g_{EI}$ | $0.25 \text{ mS}/\text{cm}^2$ |
| $g_{II}$ | $0.25 \text{ mS}/\text{cm}^2$ |
| $p_{EI}$ | 0.5 |
| $p_{IE}$ | 0.5 |
| $p_{II}$ | 0.5 |
| $\tau_{r,E}$ | 0.5 ms |
| $\tau_{\text{peak},E}$ | 0.5 ms |
| $\tau_{d,E}$ | 3 ms |
| $V_{\text{rev},E}$ | 0 mV |
| $\tau_{r,I}$ | 0.5 ms |
| $\tau_{\text{peak},I}$ | 0.5 ms |
| $\tau_{d,I}$ | 9 ms |
| $V_{\text{rev},I}$ | -75 mV |

### PING+M model

In order to reproduce the experimentally observed hysteresis effects, we implemented spike frequency adaption in the model pyramidal neurons. The PING+M model is taken from the Adaptation-based, Deterministic Weak PING model from Börgers (Chapter 32 of (Börgers, 2017)). In this model, the previous PING model is modified by the addition of a model M-Current to the pyramidal neurons. Otherwise, the network is identical to the PING model described above.

PING+M Neuron parameters:

|  |  |
| --- | --- |
| $g_M$ | 0.4 |

### LIF model

In order to investigate the generality of the model results, we next implemented a simple network of leaky-integrate-and-fire neurons. This network was composed of 80% excitatory neurons and 20% inhibitory neurons, coupled via instantaneous synapses. Excitatory neurons were not mutually connected, while the remaining connectivity was all-to-all, with synapse magnitude randomly distributed uniformly between 0 and the respective post-synaptic-potential (PSP) value. Each neuron accumulates postsynaptic potentials until the threshold for spiking is reached. Upon spiking, each neuron transmits to its synaptic partners a post synaptic event and its potential is reset. The membrane voltage of the model LIF neurons is given by: dV/dt = − V/C + I/C, with the membrane timescale tau = R*C, where R is the input resistance of the neuron, C is the membrane capacitance, and I includes both basal and synaptic currents. We drove the network with symmetric single slow sine waves or with white noise. The dynamics of the network were evaluated numerically at a resolution of tau using the Euler method.

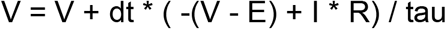

LIF Network parameters:

|  |  |
| --- | --- |
| $N_E$ | 200 |
| $N_I$ | 50 |
| $I_E$ | 1.5 nA |
| $\sigma_E$ | 0.05 $\mu A/cm^2$ |
| $I_I$ | 0 $\mu A/cm^2$ |
| $PSP_{EE}$ | 0 mV |
| $PSP_{IE}$ | 1 mV |
| $PSP_{EI}$ | -1 mV |
| $PSP_{II}$ | -1 mV |
| $\tau$ | 0.5 ms |
| $C$ | 0.5 nF |
| $R$ | 40 Mohms |
| $V_{rev,E}$ | 0 mV |
| $V_{spike}$ | 30 mV |
| $V_{\text{thresh}}$ | -40 mV |
| $V_{\text{reset}}$ | -70 mV |
| $V_{\text{leak}}$ | -60 mV |

## References

Adesnik, H., and Scanziani, M. (2010). Lateral competition for cortical space by layer-specific horizontal circuits. Nature 464, 1155–1160.

Akam, T., and Kullmann, D.M. (2010). Oscillations and filtering networks support flexible routing of information. Neuron 67, 308–320.

Akam, T., Oren, I., Mantoan, L., Ferenczi, E., and Kullmann, D.M. (2012). Oscillatory dynamics in the hippocampus support dentate gyrus-CA3 coupling. Nature neuroscience 15, 763–768.

Azouz, R., and Gray, C.M. (2003). Adaptive coincidence detection and dynamic gain control in visual cortical neurons in vivo. Neuron 37, 513–523.

Azouz, R., and Gray, C.M. (2008). Stimulus-selective spiking is driven by the relative timing of synchronous excitation and disinhibition in cat striate neurons in vivo. The European journal of neuroscience 28, 1286–1300.

Bastos, A.M., Vezoli, J., Bosman, C.A., Schoffelen, J.M., Oostenveld, R., Dowdall, J.R., De Weerd, P., Kennedy, H., and Fries, P. (2015). Visual areas exert feedforward and feedback influences through distinct frequency channels. Neuron 85, 390–401.

Besserve, M., Lowe, S.C., Logothetis, N.K., Schölkopf, B., and Panzeri, S. (2015). Shifts of Gamma Phase across Primary Visual Cortical Sites Reflect Dynamic Stimulus-Modulated Information Transfer. PLoS biology 13, e1002257.

Börgers, C. (2017). An introduction to modeling neuronal dynamics (Springer).

Börgers, C., Epstein, S., and Kopell, N.J. (2005). Background gamma rhythmicity and attention in cortical local circuits: a computational study. Proceedings of the National Academy of Sciences of the United States of America 102, 7002–7007.

Börgers, C., and Kopell, N. (2003). Synchronization in networks of excitatory and inhibitory neurons with sparse, random connectivity. Neural computation 15, 509–538.

Börgers, C., and Kopell, N.J. (2008). Gamma oscillations and stimulus selection. Neural computation 20, 383–414.

Bosman, C.A., Schoffelen, J.M., Brunet, N., Oostenveld, R., Bastos, A.M., Womelsdorf, T., Rubehn, B., Stieglitz, T., De Weerd, P., and Fries, P. (2012). Attentional stimulus selection through selective synchronization between monkey visual areas. Neuron 75, 875–888.

Boyden, E.S., Zhang, F., Bamberg, E., Nagel, G., and Deisseroth, K. (2005). Millisecond-timescale, genetically targeted optical control of neural activity. Nature neuroscience 8, 1263–1268.

Brainard, D.H. (1997). The Psychophysics Toolbox. Spat Vis 10, 433–436.

Brown, P., Salenius, S., Rothwell, J.C., and Hari, R. (1998). Cortical correlate of the Piper rhythm in humans. Journal of neurophysiology 80, 2911–2917.

Brunet, N., Bosman, C.A., Roberts, M., Oostenveld, R., Womelsdorf, T., De Weerd, P., and Fries, P. (2015). Visual cortical gamma-band activity during free viewing of natural images. Cereb Cortex 25, 918–926.

Bryant, H.L., and Segundo, J.P. (1976). Spike initiation by transmembrane current: a white-noise analysis. The Journal of physiology 260, 279–314.

Buffalo, E.A., Fries, P., Landman, R., Buschman, T.J., and Desimone, R. (2011). Laminar differences in gamma and alpha coherence in the ventral stream. Proceedings of the National Academy of Sciences of the United States of America 108, 11262–11267.

Buracas, G.T., Zador, A.M., DeWeese, M.R., and Albright, T.D. (1998). Efficient discrimination of temporal patterns by motion-sensitive neurons in primate visual cortex. Neuron 20, 959–969.

Butler, J.L., Mendonça, P.R., Robinson, H.P., and Paulsen, O. (2016). Intrinsic Cornu Ammonis Area 1 Theta-Nested Gamma Oscillations Induced by Optogenetic Theta Frequency Stimulation. The Journal of neuroscience : the official journal of the Society for Neuroscience 36, 4155–4169.

Buzsáki, G., and Wang, X.J. (2012). Mechanisms of gamma oscillations. Annual review of neuroscience 35, 203–225.

Cardin, J.A., Carlén, M., Meletis, K., Knoblich, U., Zhang, F., Deisseroth, K., Tsai, L.H., and Moore, C.I. (2009). Driving fast-spiking cells induces gamma rhythm and controls sensory responses. Nature 459, 663–667.

Chichilnisky, E.J. (2001). A simple white noise analysis of neuronal light responses. Network 12, 199–213.

Christensen, R.K., Lindén, H., Nakamura, M., and Barkat, T.R. (2019). White Noise Background Improves Tone Discrimination by Suppressing Cortical Tuning Curves. Cell reports 29, 2041–2053 e2044.

Csicsvari, J., Jamieson, B., Wise, K.D., and Buzsáki, G. (2003). Mechanisms of gamma oscillations in the hippocampus of the behaving rat. Neuron 37, 311–322.

Dhamala, M., Rangarajan, G., and Ding, M. (2008). Estimating Granger causality from fourier and wavelet transforms of time series data. Physical review letters 100, 018701.

Diester, I., Kaufman, M.T., Mogri, M., Pashaie, R., Goo, W., Yizhar, O., Ramakrishnan, C., Deisseroth, K., and Shenoy, K.V. (2011). An optogenetic toolbox designed for primates. Nature neuroscience 14, 387–397.

Douglas, R.J., and Martin, K.A. (2004). Neuronal circuits of the neocortex. Annual review of neuroscience 27, 419–451.

Engel, A.K., Fries, P., and Singer, W. (2001). Dynamic predictions: oscillations and synchrony in top-down processing. Nature reviews Neuroscience 2, 704–716.

Etter, G., van der Veldt, S., Manseau, F., Zarrinkoub, I., Trillaud-Doppia, E., and Williams, S. (2019). Optogenetic gamma stimulation rescues memory impairments in an Alzheimer’s disease mouse model. Nature communications 10, 5322.

Fellous, J.M., Houweling, A.R., Modi, R.H., Rao, R.P., Tiesinga, P.H., and Sejnowski, T.J. (2001). Frequency dependence of spike timing reliability in cortical pyramidal cells and interneurons. Journal of neurophysiology 85, 1782–1787.

Fellous, J.M., and Sejnowski, T.J. (2000). Cholinergic induction of oscillations in the hippocampal slice in the slow (0.5-2 Hz), theta (5-12 Hz), and gamma (35-70 Hz) bands. Hippocampus 10, 187–197.

Fisahn, A., Pike, F.G., Buhl, E.H., and Paulsen, O. (1998). Cholinergic induction of network oscillations at 40 Hz in the hippocampus in vitro. Nature 394, 186–189.

Fries, P. (2005). A mechanism for cognitive dynamics: neuronal communication through neuronal coherence. Trends in cognitive sciences 9, 474–480.

Fries, P. (2009). Neuronal gamma-band synchronization as a fundamental process in cortical computation. Annual review of neuroscience 32, 209–224.

Fries, P. (2015). Rhythms for Cognition: Communication through Coherence. Neuron 88, 220–235.

Fries, P., Roelfsema, P.R., Engel, A.K., König, P., and Singer, W. (1997). Synchronization of oscillatory responses in visual cortex correlates with perception in interocular rivalry. Proceedings of the National Academy of Sciences of the United States of America 94, 12699–12704.

Fries, P., Scheeringa, R., and Oostenveld, R. (2008a). Finding gamma. Neuron 58, 303–305.

Fries, P., Schröder, J.H., Roelfsema, P.R., Singer, W., and Engel, A.K. (2002). Oscillatory neuronal synchronization in primary visual cortex as a correlate of stimulus selection. The Journal of neuroscience : the official journal of the Society for Neuroscience 22, 3739–3754.

Fries, P., Womelsdorf, T., Oostenveld, R., and Desimone, R. (2008b). The effects of visual stimulation and selective visual attention on rhythmic neuronal synchronization in macaque area V4. The Journal of neuroscience : the official journal of the Society for Neuroscience 28, 4823–4835.

Gerits, A., Vancraeyenest, P., Vreysen, S., Laramee, M.E., Michiels, A., Gijsbers, R., Van den Haute, C., Moons, L., Debyser, Z., Baekelandt, V., et al. (2015). Serotype-dependent transduction efficiencies of recombinant adeno-associated viral vectors in monkey neocortex. Neurophotonics 2, 031209.

Gray, C.M., Engel, A.K., König, P., and Singer, W. (1992). Synchronization of oscillatory neuronal responses in cat striate cortex: temporal properties. Visual neuroscience 8, 337–347.

Gray, C.M., König, P., Engel, A.K., and Singer, W. (1989). Oscillatory responses in cat visual cortex exhibit inter-columnar synchronization which reflects global stimulus properties. Nature 338, 334–337.

Gray, C.M., and McCormick, D.A. (1996). Chattering cells: superficial pyramidal neurons contributing to the generation of synchronous oscillations in the visual cortex. Science 274, 109–113.

Gray, C.M., and Singer, W. (1989). Stimulus-specific neuronal oscillations in orientation columns of cat visual cortex. Proceedings of the National Academy of Sciences of the United States of America 86, 1698–1702.

Gray, C.M., and Viana Di Prisco, G. (1997). Stimulus-dependent neuronal oscillations and local synchronization in striate cortex of the alert cat. The Journal of neuroscience : the official journal of the Society for Neuroscience 17, 3239–3253.

Gregoriou, G.G., Gotts, S.J., Zhou, H., and Desimone, R. (2009). High-frequency, long-range coupling between prefrontal and visual cortex during attention. Science 324, 1207–1210.

Grothe, I., Neitzel, S.D., Mandon, S., and Kreiter, A.K. (2012). Switching neuronal inputs by differential modulations of gamma-band phase-coherence. The Journal of neuroscience : the official journal of the Society for Neuroscience 32, 16172–16180.

Grothe, I., Rotermund, D., Neitzel, S.D., Mandon, S., Ernst, U.A., Kreiter, A.K., and Pawelzik, K.R. (2018). Attention Selectively Gates Afferent Signal Transmission to Area V4. The Journal of neuroscience : the official journal of the Society for Neuroscience 38, 3441–3452.

Gulbinaite, R., Roozendaal, D.H.M., and VanRullen, R. (2019). Attention differentially modulates the amplitude of resonance frequencies in the visual cortex. NeuroImage 203, 116146.

Hadjipapas, A., Lowet, E., Roberts, M.J., Peter, A., and De Weerd, P. (2015). Parametric variation of gamma frequency and power with luminance contrast: A comparative study of human MEG and monkey LFP and spike responses. NeuroImage 112, 327–340.

Hahn, G., Bujan, A.F., Frégnac, Y., Aertsen, A., and Kumar, A. (2014). Communication through resonance in spiking neuronal networks. PLoS computational biology 10, e1003811.

Harris, K.D., Csicsvari, J., Hirase, H., Dragoi, G., and Buzsáki, G. (2003). Organization of cell assemblies in the hippocampus. Nature 424, 552–556.

Hasenstaub, A., Shu, Y., Haider, B., Kraushaar, U., Duque, A., and McCormick, D.A. (2005). Inhibitory postsynaptic potentials carry synchronized frequency information in active cortical networks. Neuron 47, 423–435.

Hutcheon, B., and Yarom, Y. (2000). Resonance, oscillation and the intrinsic frequency preferences of neurons. Trends in neurosciences 23, 216–222.

Iaccarino, H.F., Singer, A.C., Martorell, A.J., Rudenko, A., Gao, F., Gillingham, T.Z., Mathys, H., Seo, J., Kritskiy, O., Abdurrob, F., et al. (2016). Gamma frequency entrainment attenuates amyloid load and modifies microglia. Nature 540, 230–235.

Jia, X., Xing, D., and Kohn, A. (2013). No consistent relationship between gamma power and peak frequency in macaque primary visual cortex. The Journal of neuroscience : the official journal of the Society for Neuroscience 33, 17–25.

Kruse, W., and Eckhorn, R. (1996). Inhibition of sustained gamma oscillations (35-80 Hz) by fast transient responses in cat visual cortex. Proceedings of the National Academy of Sciences of the United States of America 93, 6112–6117.

Lampl, I., and Yarom, Y. (1997). Subthreshold oscillations and resonant behavior: two manifestations of the same mechanism. Neuroscience 78, 325–341.

Lee, J.H., Whittington, M.A., and Kopell, N.J. (2013). Top-down beta rhythms support selective attention via interlaminar interaction: a model. PLoS computational biology 9, e1003164.

Lowet, E., Roberts, M.J., Peter, A., Gips, B., and De Weerd, P. (2017). A quantitative theory of gamma synchronization in macaque V1. eLife 6.

Lu, Y., Truccolo, W., Wagner, F.B., Vargas-Irwin, C.E., Ozden, I., Zimmermann, J.B., May, T., Agha, N.S., Wang, J., and Nurmikko, A.V. (2015). Optogenetically induced spatiotemporal gamma oscillations and neuronal spiking activity in primate motor cortex. Journal of neurophysiology 113, 3574–3587.

Mainen, Z.F., and Sejnowski, T.J. (1995). Reliability of spike timing in neocortical neurons. Science 268, 1503–1506.

Maris, E., and Oostenveld, R. (2007). Nonparametric statistical testing of EEG- and MEG-data. Journal of neuroscience methods 164, 177–190.

Marmarelis, P.Z., and Naka, K. (1972). White-noise analysis of a neuron chain: an application of the Wiener theory. Science 175, 1276–1278.

Michalareas, G., Vezoli, J., van Pelt, S., Schoffelen, J.M., Kennedy, H., and Fries, P. (2016). Alpha-Beta and Gamma Rhythms Subserve Feedback and Feedforward Influences among Human Visual Cortical Areas. Neuron 89, 384–397.

Mitchell, J.F., Sundberg, K.A., and Reynolds, J.H. (2009). Spatial attention decorrelates intrinsic activity fluctuations in macaque area V4. Neuron 63, 879–888.

Munk, M.H., Roelfsema, P.R., König, P., Engel, A.K., and Singer, W. (1996). Role of reticular activation in the modulation of intracortical synchronization. Science 272, 271–274.

Ni, J., Wunderle, T., Lewis, C.M., Desimone, R., Diester, I., and Fries, P. (2016). Gamma-Rhythmic Gain Modulation. Neuron 92, 240–251.

Nichols, T.E., and Holmes, A.P. (2002). Nonparametric permutation tests for functional neuroimaging: a primer with examples. Hum Brain Mapp 15, 1–25.

Onorato, I., Neuenschwander, S., Hoy, J., Lima, B., Rocha, K.S., Broggini, A.C., Uran, C., Spyropoulos, G., Klon-Lipok, J., Womelsdorf, T., et al. (2020). A Distinct Class of Bursting Neurons with Strong Gamma Synchronization and Stimulus Selectivity in Monkey V1. Neuron 105, 180–197 e185.

Oostenveld, R., Fries, P., Maris, E., and Schoffelen, J.M. (2011). FieldTrip: Open source software for advanced analysis of MEG, EEG, and invasive electrophysiological data. Computational intelligence and neuroscience 2011, 156869.

Palmigiano, A., Geisel, T., Wolf, F., and Battaglia, D. (2017). Flexible information routing by transient synchrony. Nature neuroscience 20, 1014–1022.

Payne, B.R. (1993). Evidence for visual cortical area homologs in cat and macaque monkey. Cereb Cortex 3, 1–25.

Pesaran, B., Pezaris, J.S., Sahani, M., Mitra, P.P., and Andersen, R.A. (2002). Temporal structure in neuronal activity during working memory in macaque parietal cortex. Nature neuroscience 5, 805–811.

Pesaran, B., Vinck, M., Einevoll, G.T., Sirota, A., Fries, P., Siegel, M., Truccolo, W., Schroeder, C.E., and Srinivasan, R. (2018). Investigating large-scale brain dynamics using field potential recordings: analysis and interpretation. Nature neuroscience 21, 903–919.

Pillow, J.W., Shlens, J., Paninski, L., Sher, A., Litke, A.M., Chichilnisky, E.J., and Simoncelli, E.P. (2008). Spatio-temporal correlations and visual signalling in a complete neuronal population. Nature 454, 995–999.

Ray, S., and Maunsell, J.H. (2010). Differences in gamma frequencies across visual cortex restrict their possible use in computation. Neuron 67, 885–896.

Roberts, M.J., Lowet, E., Brunet, N.M., Ter Wal, M., Tiesinga, P., Fries, P., and De Weerd, P. (2013). Robust gamma coherence between macaque V1 and V2 by dynamic frequency matching. Neuron 78, 523–536.

Rodriguez, R., Kallenbach, U., Singer, W., and Munk, M.H. (2004). Short- and long-term effects of cholinergic modulation on gamma oscillations and response synchronization in the visual cortex. The Journal of neuroscience : the official journal of the Society for Neuroscience 24, 10369–10378.

Rohenkohl, G., Bosman, C.A., and Fries, P. (2018). Gamma Synchronization between V1 and V4 Improves Behavioral Performance. Neuron 100, 953–963 e953.

Salinas, E., and Sejnowski, T.J. (2001). Correlated neuronal activity and the flow of neural information. Nature reviews Neuroscience 2, 539–550.

Scheyltjens, I., Laramee, M.E., Van den Haute, C., Gijsbers, R., Debyser, Z., Baekelandt, V., Vreysen, S., and Arckens, L. (2015). Evaluation of the expression pattern of rAAV2/1, 2/5, 2/7, 2/8, and 2/9 serotypes with different promoters in the mouse visual cortex. J Comp Neurol 523, 2019–2042.

Schreiber, S., Fellous, J.M., Tiesinga, P., and Sejnowski, T.J. (2004). Influence of ionic conductances on spike timing reliability of cortical neurons for suprathreshold rhythmic inputs. Journal of neurophysiology 91, 194–205.

Sherfey, J.S., Ardid, S., Hass, J., Hasselmo, M.E., and Kopell, N.J. (2018). Flexible resonance in prefrontal networks with strong feedback inhibition. PLoS computational biology 14, e1006357.

Sohal, V.S., Zhang, F., Yizhar, O., and Deisseroth, K. (2009). Parvalbumin neurons and gamma rhythms enhance cortical circuit performance. Nature 459, 698–702.

Stark, E., Roux, L., Eichler, R., Senzai, Y., Royer, S., and Buzsáki, G. (2014). Pyramidal cell-interneuron interactions underlie hippocampal ripple oscillations. Neuron 83, 467–480.

Tchumatchenko, T., Newman, J.P., Fong, M.F., and Potter, S.M. (2013). Delivery of continuously-varying stimuli using channelrhodopsin-2. Frontiers in neural circuits 7, 184.

Tiesinga, P., and Sejnowski, T.J. (2009). Cortical enlightenment: are attentional gamma oscillations driven by ING or PING? Neuron 63, 727–732.

Traub, R.D., Whittington, M.A., Colling, S.B., Buzsáki, G., and Jefferys, J.G. (1996). Analysis of gamma rhythms in the rat hippocampus in vitro and in vivo. The Journal of physiology 493 (Pt 2), 471–484.

Traub, R.D., Wong, R.K., Miles, R., and Michelson, H. (1991). A model of a CA3 hippocampal pyramidal neuron incorporating voltage-clamp data on intrinsic conductances. Journal of neurophysiology 66, 635–650.

van Kerkoerle, T., Self, M.W., Dagnino, B., Gariel-Mathis, M.A., Poort, J., van der Togt, C., and Roelfsema, P.R. (2014). Alpha and gamma oscillations characterize feedback and feedforward processing in monkey visual cortex. Proceedings of the National Academy of Sciences of the United States of America.

VanRullen, R., and Macdonald, J.S. (2012). Perceptual echoes at 10 Hz in the human brain. Current biology : CB 22, 995–999.

Varela, F., Lachaux, J.P., Rodriguez, E., and Martinerie, J. (2001). The brainweb: phase synchronization and large-scale integration. Nature reviews Neuroscience 2, 229–239.

Vasileva, A., and Jessberger, R. (2005). Precise hit: adeno-associated virus in gene targeting. Nat Rev Microbiol 3, 837–847.

Vinck, M., van Wingerden, M., Womelsdorf, T., Fries, P., and Pennartz, C.M. (2010). The pairwise phase consistency: A bias-free measure of rhythmic neuronal synchronization. NeuroImage 51, 112–122.

Vite, C.H., Passini, M.A., Haskins, M.E., and Wolfe, J.H. (2003). Adeno-associated virus vector-mediated transduction in the cat brain. Gene Ther 10, 1874–1881.

Wang, X.J. (1999). Synaptic basis of cortical persistent activity: the importance of NMDA receptors to working memory. The Journal of neuroscience : the official journal of the Society for Neuroscience 19, 9587–9603.

Wang, X.J. (2010). Neurophysiological and computational principles of cortical rhythms in cognition. Physiological reviews 90, 1195–1268.

Wang, X.J., and Buzsáki, G. (1996). Gamma oscillation by synaptic inhibition in a hippocampal interneuronal network model. The Journal of neuroscience : the official journal of the Society for Neuroscience 16, 6402–6413.

Whittington, M.A., and Traub, R.D. (2003). Interneuron diversity series: inhibitory interneurons and network oscillations in vitro. Trends in neurosciences 26, 676–682.

Wilson, H.R., and Cowan, J.D. (1972). Excitatory and inhibitory interactions in localized populations of model neurons. Biophysical journal 12, 1–24.

Womelsdorf, T., Lima, B., Vinck, M., Oostenveld, R., Singer, W., Neuenschwander, S., and Fries, P. (2012). Orientation selectivity and noise correlation in awake monkey area V1 are modulated by the gamma cycle. Proceedings of the National Academy of Sciences of the United States of America 109, 4302–4307.

Womelsdorf, T., Schoffelen, J.M., Oostenveld, R., Singer, W., Desimone, R., Engel, A.K., and Fries, P. (2007). Modulation of neuronal interactions through neuronal synchronization. Science 316, 1609–1612.

