## Supplemental figures and text for "Cortical resonance selects coherent input"

Supplemental information consists of :

Supplemental Figure S1, related to Figure 1

Supplemental Figure S2, related to Figure 1

Supplemental Figure S3, related to Figure 2

Supplemental Figure S4, related to Figure 2

Supplemental Figure S5, related to Figure 3

Supplemental Figure S6, related to Figure 4

Supplemental Figure S7, related to Figure 5

Supplemental Text

#### Supplemental Figure Legends

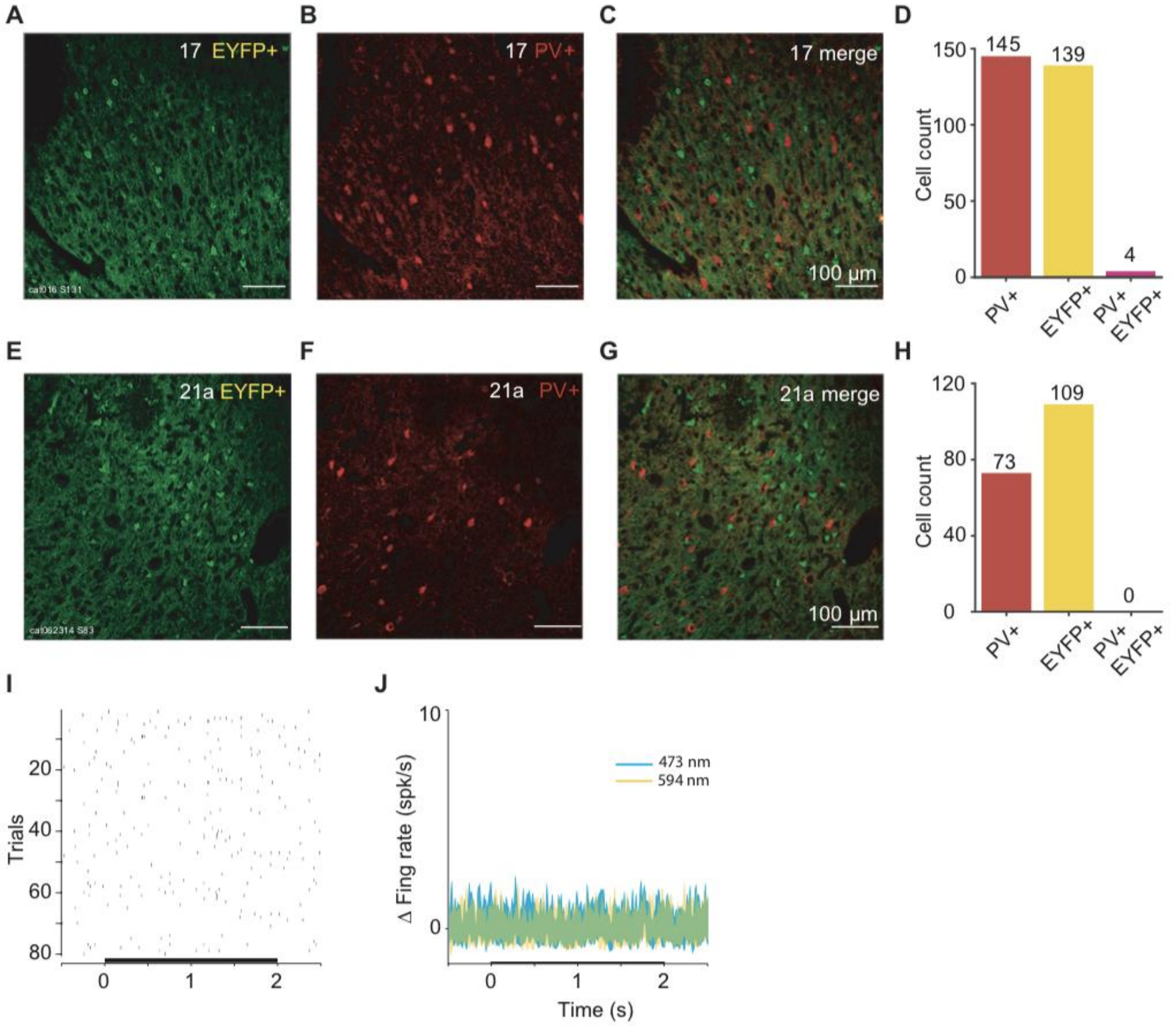


**Supplemental Figure S1. Viral transfection was largely selective for excitatory neurons.** (A-C) Confocal images of immunohistochemistry performed on slices from area 17 after viral transfection. (A) Endogenous fluorescence of ChR2-eYFP. (B) Fluorescence of secondary antibody after staining for PV+. (C) Merged images, testing for neuronal co-labeling with ChR2-eYFP and PV+ antibody. (D) Counts of PV+ labeled neurons, EYFP+ labeled neurons, and co-labeled neurons in area 17. (E-H) Same as A-D, but for area 21a. No co-labeled neurons can be found. (I) Example MUA response to 2 s of blue laser stimulation to a site in area 21a not expressing ChR2. (J) Group results showing firing rate changes from baseline for 2 s of constant blue and yellow laser stimulation for 8 sites (5 area 21a, 3 area 17) in 4 cats.

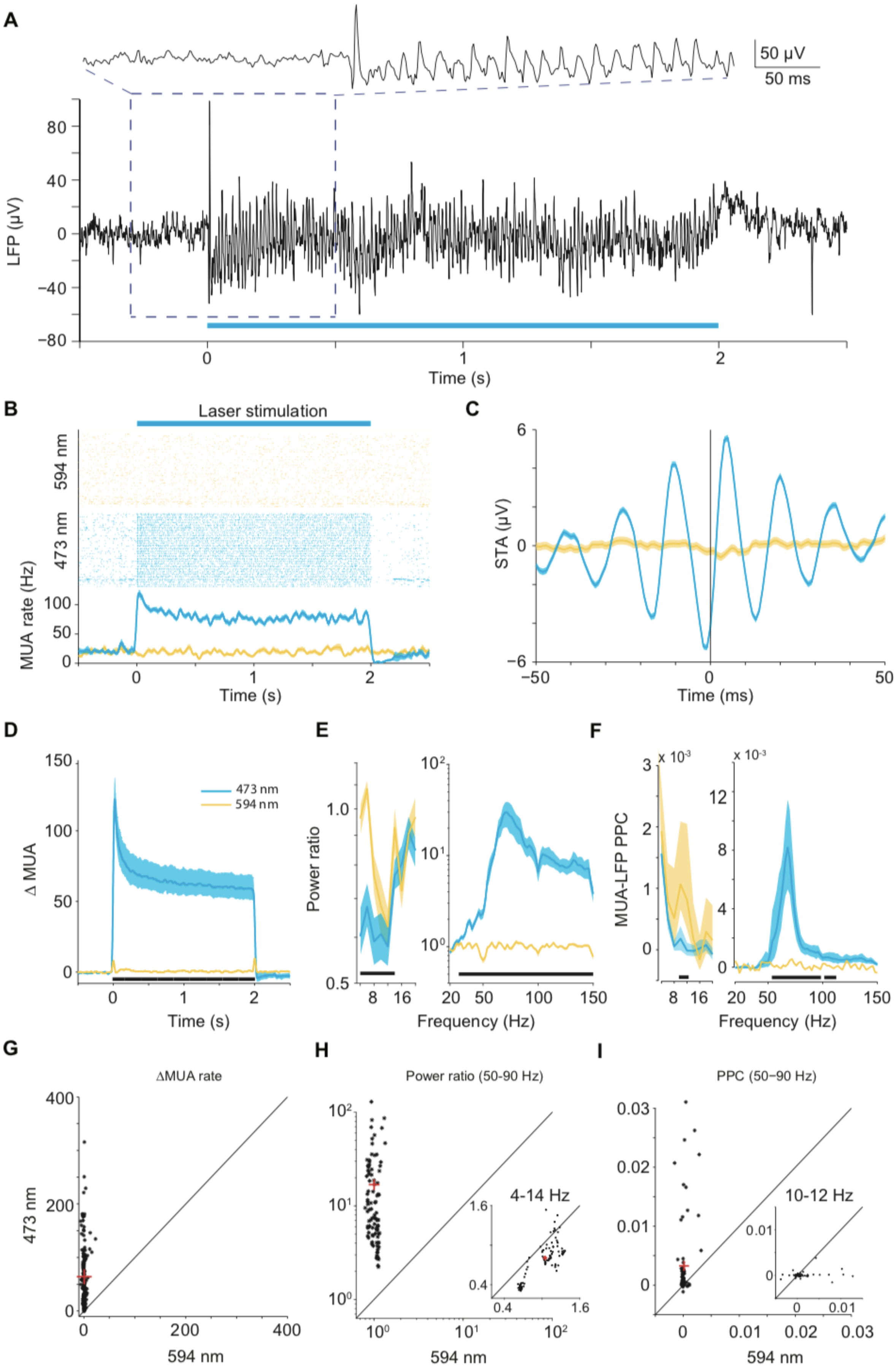

A
50 µV
50 ms
LFP (µV)
80
40
0
−40
−80
0
1
2
Time (s)
B
Laser stimulation
594 nm
473 nm
MUA rate (Hz)
100
50
0
0
1
2
Time (s)
C
6
STA (µV)
0
−6
−50
0
50
Time (ms)
D
150
100
50
0
Δ MUA
473 nm
594 nm
0
1
2
Time (s)
E
1.0
0.5
Power ratio
8
16
10²
10¹
10⁰
20
50
100
150
Frequency (Hz)
F
x 10⁻³
3
2
1
0
MUA-LFP PPC
8
16
x 10⁻³
12
8
4
0
20
50
100
150
Frequency (Hz)
G
ΔMUA rate
400
200
0
473 nm
0
200
400
594 nm
H
Power ratio (50-90 Hz)
10²
10¹
10⁰
4-14 Hz
1.6
0.4
0.4
1.6
10⁰
10¹
10²
594 nm
I
PPC (50−90 Hz)
0.03
0.02
0.01
0
10-12 Hz
0.01
0
0
0.01
0
0.01
0.02
0.03
594 nm

**Supplemental Figure S2. Group results for MUA rate, LFP power and MUA-LFP PPC induced by constant light.** (A) Example recording site in area 21a shows strong gamma-band synchronization in the local field potential induced by constant-illumination. (B) Robust MUA response to constant illumination at the same site. Blue: 473 nm wavelength light; Yellow: 594 nm wavelength light. Shaded area indicates $\pm 1$ SEM. (C) Spike-triggered LFP for example data shown in A and B. Shaded area indicates $\pm 1$ SEM. (D) Average MUA spike density change due to optogenetic stimulation. Smoothed by a Gaussian function ($\sigma$ = 12.5 ms, truncated at $\pm 2\sigma$). (E) Average LFP power ratio (optogenetic stimulation versus baseline) spectrum. Note different y-axis scales for lower and higher frequency ranges. (F) Average MUA-LFP PPC spectrum. Note different y-axis scales for lower and higher frequency ranges. (E, F) use ±0.5 s epochs for the analyses from 4 to 20 Hz, and ±0.25 s long epochs for the analyses from 20-150 Hz. (B-F) Blue (yellow) lines show data obtained with 473 nm (594 nm) light stimulation. Shaded areas indicate $\pm 1$ SEM across recording sites, which is shown for illustration only. Black bars at the bottom indicate frequency ranges with statistically significant ($p < 0.05$) differences between blue and yellow light stimulation, based on a cluster-level permutation test including correction for the multiple comparisons across frequencies. For the main clusters from panels (D-F), panels (G-I) illustrate the underlying distributions as scatter plots. (G) Each dot shows the MUA rate (0.3-2 s after light onset) of one recording site for blue light on the y-axis versus yellow light at the x-axis. The red cross corresponds to the respective median values. (H) Same as (G), but for the LFP power ratio. The main plot is for the gamma band (50-90 Hz); the inset plot for the low-frequency cluster from (E) (4-14 Hz). (I) Same as (G), but for MUA-LFP PPC during light stimulation. Each dot corresponds to one MUA recording site. The main plot is for the gamma band (50-90 Hz); the inset plot is for the low-frequency cluster from (F) (10-12 Hz).

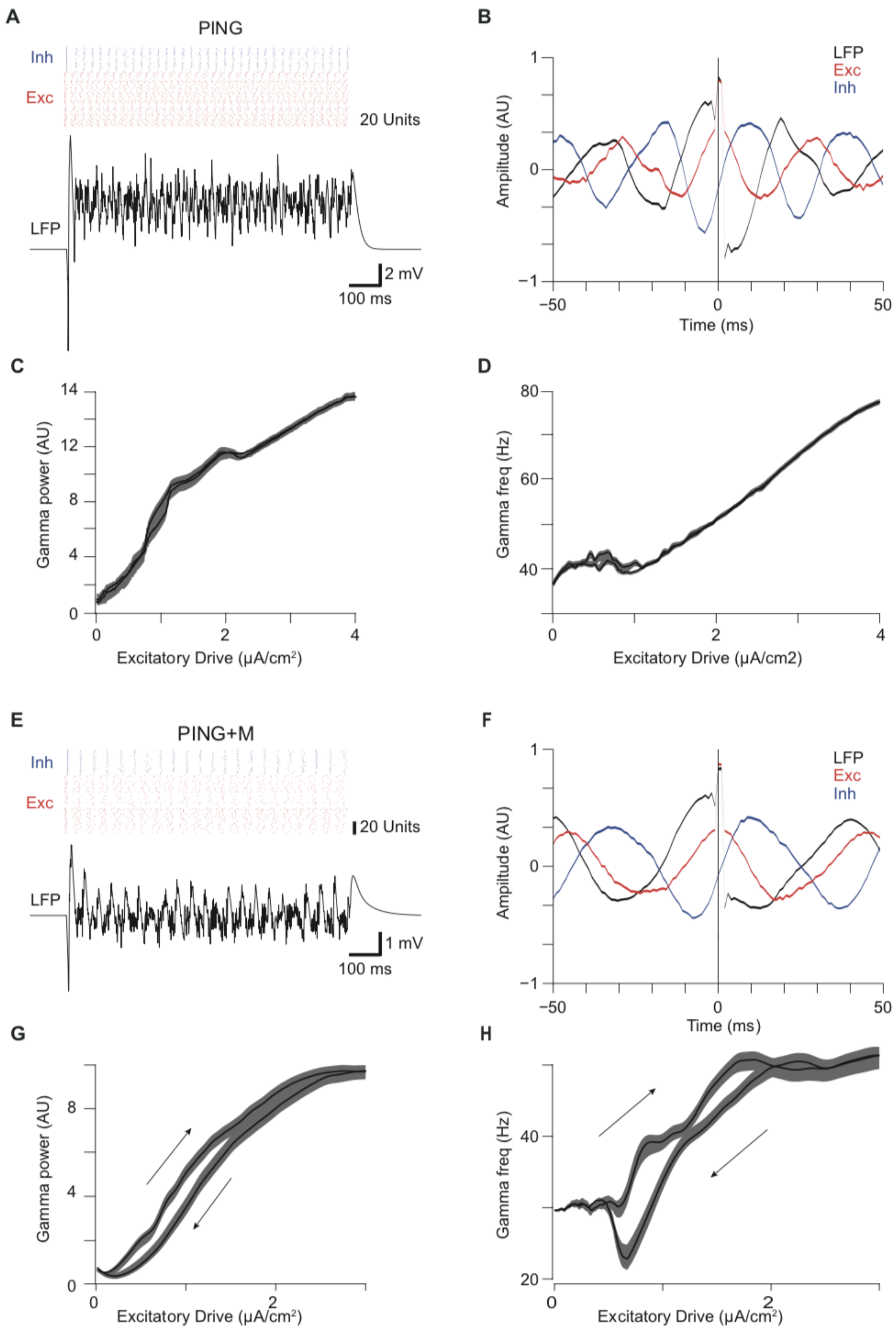


**Supplemental Figure S3. Synchronization in the PING and PING+M models.** (A-B) Response of the PING model to constant stimulation. (A) Raster plot of inhibitory (Inh) and excitatory (Exc) neuron spiking, and average membrane potential (LFP) reveal robust gamma-band synchronization in the PING model in response to constant excitation. (B) Spike-triggered averaging, based on spikes of excitatory units, in the PING model reflects characteristic gamma cycle with excitation leading inhibition. (C) Gamma power and (D) frequency increase with increasing excitatory drive in the PING network, but do not demonstrate hysteresis. (E-F) Response of the PING+M model to constant stimulation. (E) Raster plot of inhibitory (Inh) and excitatory (Exc) neuron spiking, and average membrane potential (LFP) reveal robust gamma-band synchronization in the PING+M model, but at a lower frequency as compared to the PING model, when constant excitation is matched. (F) As in (B), but for the PING+M model. (G) In the PING+M model, gamma power and (H) frequency increase with increasing excitatory drive. Arrows indicate hysteresis in response to increasing (upper arrow) versus decreasing (lower arrow) laser power, in qualitative agreement with recordings.

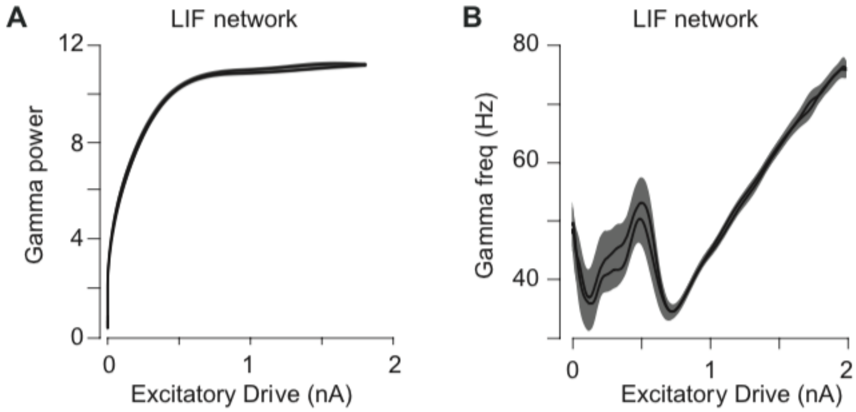


**Supplemental Figure S4. Synchronization of LIF network with increased excitatory drive.** (A-B) LIF network exhibits increased gamma power (A) and frequency (B) with increased excitatory drive, but does not display hysteresis.

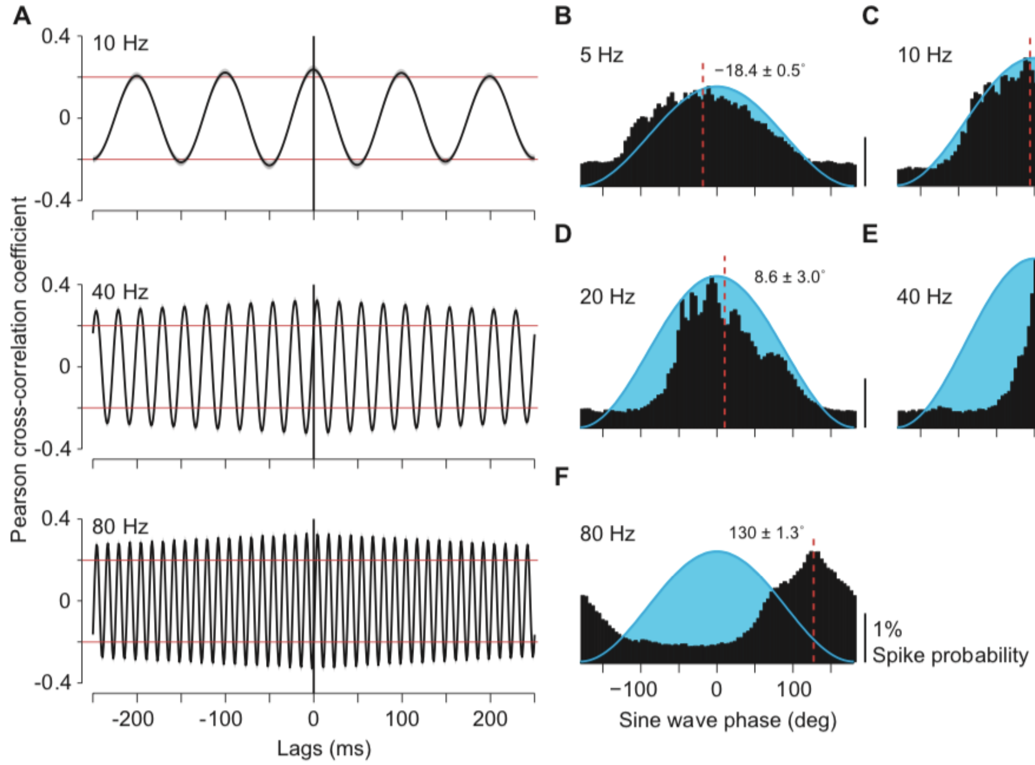


**Supplemental Figure S5. MUA responses to sinusoidal stimulation.** (A) The Pearson cross-correlation coefficients between sinusoidal optogenetic drive and MUA for stimulation at 10 Hz, 40 Hz and 80 Hz demonstrate increased correlation in the gamma-band. Red horizontal lines are shown at ±0.2 in all panels to easy comparison. (B-F) MUA spike probability, averaged over recording sites, as a function of the phase of the optogenetic sine wave stimulation. The optogenetic sine wave is indicated by the blue-shaded region. Each panel shows the data obtained with the frequency indicated on top of the panel. MUA responses were fitted with Gaussians, and the resulting peak latencies are indicated by dashed red lines. Peak latencies and their SEM (estimated through a jackknife procedure) are indicated as text insets. Latencies are expressed relative to the time of peak light intensity.

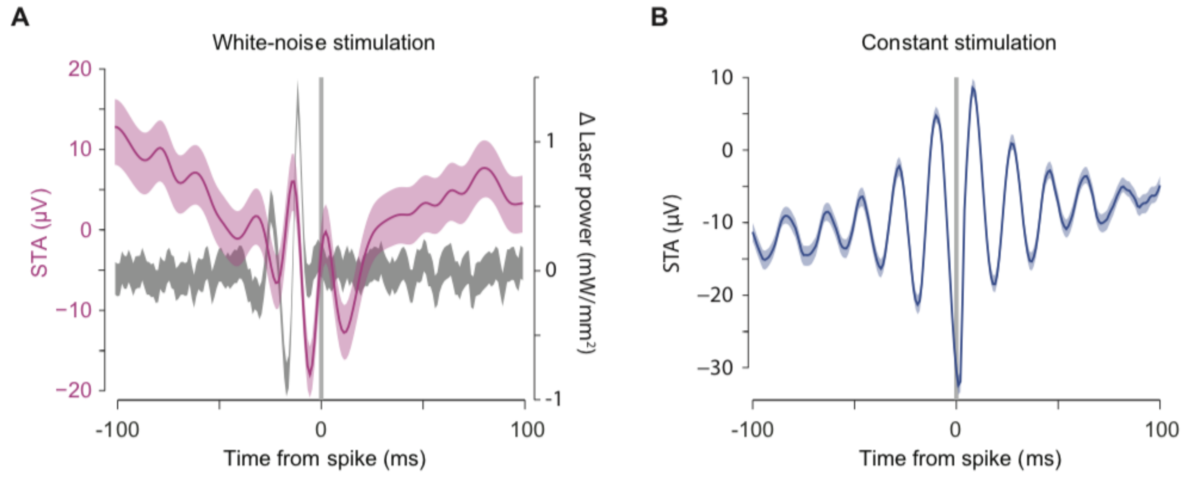


**Supplemental Figure S6. Time scale of gamma-band synchronization during white-noise and constant stimulation.** (A) Spike-triggered white noise (grey) and spike-triggered LFP (red), during white-noise stimulation. (B) Spike-triggered LFP for constant optogenetic stimulation of the same average laser power as in (A).

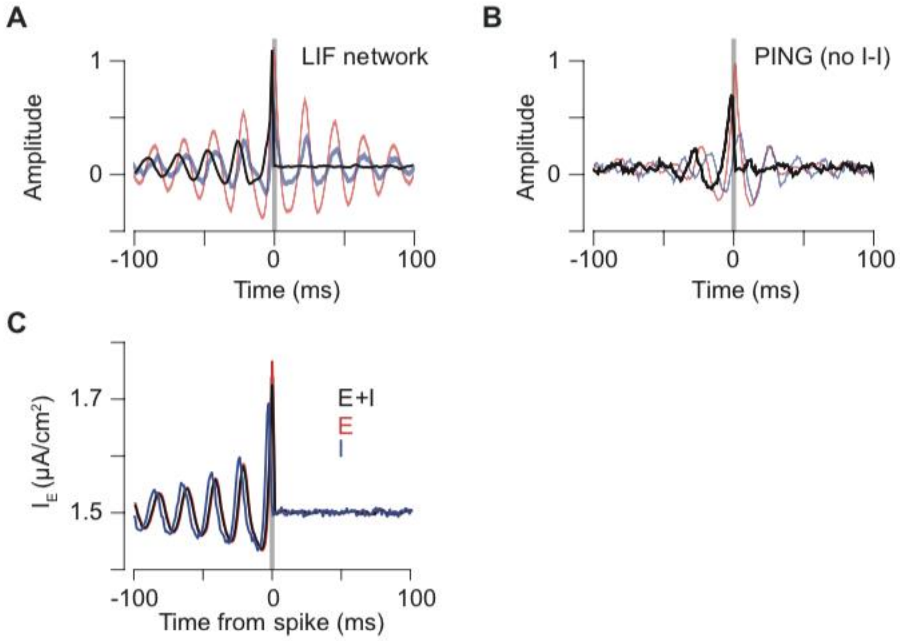


**Supplemental Figure S7. Selective transmission in LIF network and in PING network without I-I connectivity.** (A) LIF network exhibits selective transmission of coherent input. (B) PING network without I-to-I coupling exhibits selective transmission of coherent input, with reduced effect strength and lower frequency. (C) Spike-triggered-average of white-noise for different components of the PING network: all units (black), excitatory units (red), and inhibitory units (blue).

### Supplemental Text

#### Estimation of response latency with sinusoidal stimulation

Sinusoidal stimulation of different frequencies enabled estimation of neuronal response latencies. This is highly relevant when optogenetic stimulation is used to produce temporal activation patterns at high frequencies. In addition, it validates that the responses we observe are a result of optogenetic stimulation: Neuronal response latencies to optogenetic stimulation are typically on the order of 3-8 ms; By contrast, shorter latency responses are likely to reflect photo-electric artifacts (Cardin et al., 2010). To investigate response latencies, we averaged MUA responses aligned to the peaks of the sinusoids (Fig. S5B-F). During sinusoidal stimulation, the light was modulated between the respective maximal intensity and nearly zero intensity. Thus, the light crossed the threshold for effective neuronal stimulation at an unknown intensity, and it is not possible to calculate response latencies in the same way as has been done for pulse trains. Therefore, we used a technique of latency estimation that has been developed in the study of synchronized oscillations, and that is based on the slope of the spectrum of the relative phase between two signals (Schoffelen et al., 2005), in our case the light intensity and the MUA. Figure 3C shows this relative-phase spectrum and reveals a strictly linear relationship between relative phase and frequency. A linear frequency-phase relation is a signature of a fixed time lag, because a given time lag translates into increasing phase lags for increasing frequencies (Schoffelen et al., 2005). The slope of this linear relationship allowed us to infer a latency of 5.5 ms, in good agreement with previous reports of neuronal latencies.

#### References

Cardin, J.A., Carlén, M., Meletis, K., Knoblich, U., Zhang, F., Deisseroth, K., Tsai, L.H., and Moore, C.I. (2010). Targeted optogenetic stimulation and recording of neurons in vivo using cell-type-specific expression of Channelrhodopsin-2. Nature protocols 5, 247-254.

Schoffelen, J.M., Oostenveld, R., and Fries, P. (2005). Neuronal coherence as a mechanism of effective corticospinal interaction. Science 308, 111-113.
